# Actin-driven protrusions generate rapid long-range membrane tension propagation in cells

**DOI:** 10.1101/2022.09.07.507005

**Authors:** Henry De Belly, Shannon Yan, Hudson Borja da Rocha, Sacha Ichbiah, Jason P. Town, Hervé Turlier, Carlos Bustamante, Orion D. Weiner

## Abstract

Membrane tension is thought to be a long-range integrator of cell physiology. This role necessitates effective tension transmission across the cell. However, the field remains strongly divided as to whether cell membranes support or resist tension propagation, in part due to a lack of adequate tools for locally manipulating membrane tension. We overcome these limitations by leveraging optogenetics to generate localized actin-based protrusions while concurrently monitoring the propagation of membrane tension using dual-trap optical tweezers. Surprisingly, actin-driven protrusions elicit rapid global membrane tension propagation with little to no attenuation, while forces applied to the cell membrane only do not. We present a simple unifying mechanical model in which mechanical forces that act on both the membrane and actin cortex drive rapid, robust membrane tension propagation.

**Summary:** Mechanical perturbations acting on both actin cortex and plasma membrane drive global membrane tension increase within seconds

## Introduction

Membrane tension is a central regulator of many fundamental biological processes (1–9). In the context of cell migration, membrane tension has been proposed to serve as a rapid long-range coordinator of information across the cell for a ‘winner-take-all’ in polarity establishment (10–12). However, whether cell membranes support or resist long-range membrane tension propagation remains controversial. While some studies report that membrane tension rapidly propagates across the cell (10, 11, 13–15), others find little to no propagation of membrane tension (16–19). These discrepancies could stem from differences in cell types examined—for instance, membrane tension transmission is perhaps more efficient in motile cells than non-motile epithelial cells (17, 20, 21). Alternatively, the discrepancies could be due to differences in methodology; for example, exogenously-applied mechanical perturbations may elicit different tension responses than endogenously-generated mechanical forces.

## Results

To investigate membrane tension propagation upon endogenous force generation, we employed optogenetic regulation (Opto-PI3K) (22) in neutrophil-like HL-60 cells to activate localized actin-driven membrane protrusions (Fig. 1A, Fig. S1A; Fig. 1B, movie S1-3) and increase membrane tension at the protruding site (10, 13, 23). The propagation of membrane tension can be probed via a membrane tether pulled out on the opposite side of cell body using a bead (coated with lectin to bind carbohydrate groups on the membrane) and held by an optical trap (a.k.a. trap-based tether pulling assay; Fig. 1C, Fig. S2; see Methods). We observed a rapid, robust, long-range increase in membrane tension over multiple rounds of light-induced actin-driven protrusion (Fig. 1D-E, Fig. S3A-C, movie S4). The rise in tension within 5-15s of light activation is in stark contrast to the conclusion arrived at in recent studies (16), stating that “cell membranes resist flow.” We verified that the observed increase in tension correlates with the local activation of actin regulator, Rac GTPase, which is downstream of phosphoinositide 3-kinase (PI3K) activation and precedes actin-driven protrusion (Fig. S1). As a control, we treated the cells with the actin inhibitor Latrunculin B and observed a lack of membrane tension increase following light activation (Fig. 1E-F, Fig. S3D). These results demonstrate that actin-based protrusions elicit a rapid long-range propagation of membrane tension.

**Fig. 1.**
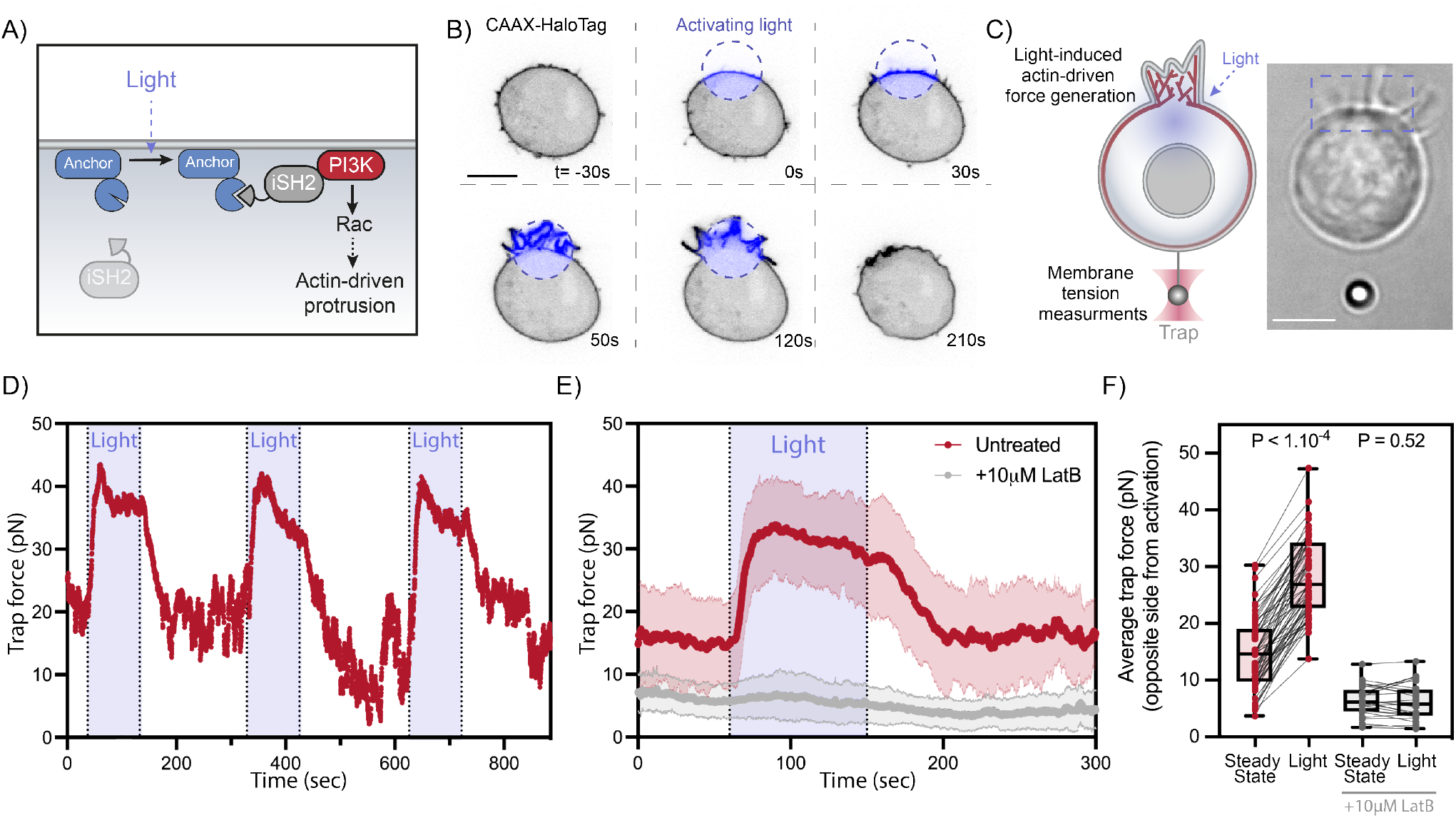
Local cell protrusions elicit a rapid long-range increase in membrane tension. (A) Optogenetic control for light-induced activation of phosphoinositide 3-kinase (PI3K) via localized recruitment of inter SH2 domain (iSH2), followed by Rac GTPase activation to initiate actin-driven cell protrusions (see Methods). (B) Time-lapse confocal images of a neutrophil-like HL-60 cell expressing opto-construct (Opto-PI3K) and membrane marker (CAAX-HaloTag) showing localized membrane protrusion upon light activation. (C) Following light-activated protrusion on one side of the cell (top), changes in membrane tension on the opposite side (bottom) are measured via a membrane tether held by an optical trap. Right, brightfield image of a protruding cell during tether pulling assay. (D) Representative time trace of trap force (a direct readout of cell membrane tension change) reveals robust and sharp increase in membrane tension over repeating cycles of light-activated protrusion on the opposite end of the cell (panel C); light: 90s on (shaded area). (E) Red: averaged time trace of trap force before (Steady-state), during (Light), and after activating cell protrusion (means ± SD; n>60, N=8). Grey: as a control, averaged trace from cells treated with actin polymerization inhibitor (10 *μ*M Latrunculin B) shows little membrane tension change upon optogenetic activation. (F) Averaged trap force before (Steady-state) and during activation. Box and whiskers: median and min to max; p values from Wilcoxon paired Student’s t test. Scale bars: 5*μ*m.

To examine the dynamics of tension propagation in more detail, we performed a dual-tether pulling assay and simultaneously monitored membrane tension on the side and the back of the cell (at 90° and 180° from the site of illumination, respectively) throughout multiple cycles of light-induced protrusion (Fig. 2A, Fig. S4; movie S5). Interestingly, the two membrane tethers exhibit a near-simultaneous increase in tension, with a delay on average of 1.2 *±* 1.2s between the two (Fig. S4E). Readouts on both tethers plateau towards similar tension levels (Fig. 2B-C). Furthermore, membrane tension measurements of the two tethers remain highly correlated during light-activated protrusion and during recovery (Fig. S4D). Our experiments indicate that endogenous actin-based protrusions generate a long-range increase in membrane tension that is transmitted virtually undampened throughout the cell within seconds.

**Fig. 2.**
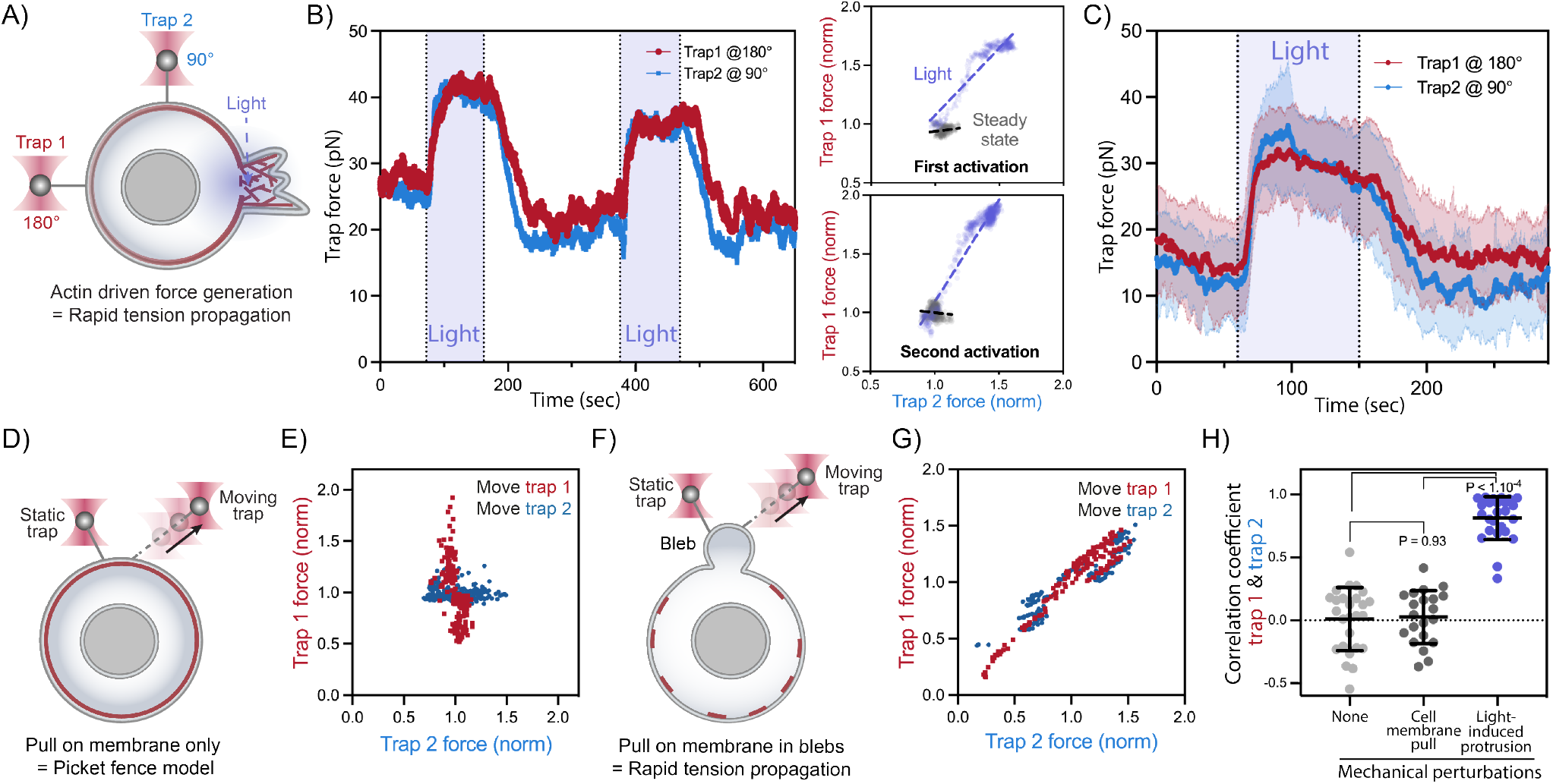
Actin-driven protrusions stimulate global membrane tension propagation, while direct pulling on the cell membrane does not. (A) A dual-tether pulling assay to simultaneously monitor membrane tension on the far-end (left, trap 1 at 180*°*) and on the side of the cell (top, trap 2 at 90°) during light-activated protrusion. (B) Representative time traces of dual trap forces over successive cycles of light-activated protrusion show coinciding tension increases on both membrane tethers adjacent to (trap 2) and at the opposite cell surface from (trap 1) protrusion; light: 90s on (shaded area), 180s off. Right: correlation between trap forces at the two tether positions during activation (blue) remains robust from first activation cycle to the next; for comparison, minimal correlation is seen between the two tethers before optogenetic activation (grey). Dashed line: linear regression. (C) Averaged traces of dual trap forces before, during (Light), and after activation (n>25, N=4). (D) A dual-tether assay to detect tension propagation (static tether, left) while a force is applied nearby by using bead to pull on membrane 2 *μ*m away (moving tether, right). (E) Trap forces (readout of membrane tension response) from the static tether remain uncorrelated to that of the moving tether (i.e., little to no change in tension on the static tether during pulling of the moving tether; Fig. S5). (F)(G) Similar to (D)(E) but probing tension in blebs (membrane detached from actin cortex); here a high correlation is observed between static and moving tethers. (H) Pearson correlation coefficient between dual trap forces measured before perturbations (None), during membrane pulling from a moving tether (panel D), and upon light-activated protrusions. Error bar: means ± SD; p values from Welch’s unpaired Student’s t test (n>20, N>3).

The contradictory observations between our current work and previous studies (16, 17, 20, 21) may originate from how a mechanical perturbation is applied to cell membranes. Here, we optogenetically induce cellular membrane protrusion (i.e., endogenous actin-driven), eliciting rapid global membrane tension propagation. In our approach, forces are potentially applied to both the actin cortex and the plasma membrane. In contrast, previous studies concluding that membrane tension is locally constrained by the actin cytoskeleton (16) used a pair of membrane tethers to pull on the cell membrane (i.e., exogenous bead-pulling), thereby applying forces to the plasma membrane alone. To test whether membrane tether-induced forces also fail to propagate in our cells, we repurposed our dual-tether assay to pull on membrane tether by actively moving one trap while measuring membrane tension on a nearby membrane tether held in place by the second trap (i.e., Fig. 2A vs. Fig. 2D). In line with analogous experiments performed in epithelial cells, we observe no propagation of membrane tension from the extending tether to the static one (Fig. 2E; Fig. S5A-B; movie S6)—even with the two tethers in close proximity (<2*μ*m apart). In contrast, when we performed the same dual-tether assay on cellular blebs (membrane detached from actin cortex), tension propagates almost instantly (<100*μ*s, i.e., below the temporal resolution of the tweezers instrument; Fig. 2F-G, Fig. S5E-J ; movie S7), in agreement with similar measurements in epithelial cells (16). Our observations that effective membrane tension propagation depends on how mechanical perturbations are exerted to the cell suggest that existing disagreements in the field are at least partially methodological in nature. While mechanical perturbations via exogenous tether pulling fail to elicit membrane tension propagation (consistent with the ‘picket fence’ model (16, 21)), endogenous actin-based force generation efficiently promotes membrane tension propagation across the cell.

Next, we investigated the mechanism of tension propagation from the site of protrusion to the rest of the cell. We observed plasma membrane enrichment in the optogenetically-induced protrusions(Fig. 1B; movie S8) on a similar timescale to the cellular deformation and tension increase (10-15s) following light activation (Fig. S1C-D). We hypothesized that membrane and cortical flows could underlie the rapid propagation of membrane tension from the protrusion to the rest of the cell. To resolve the time-dependent flow of the plasma membrane and actin cytoskeleton relative to light-activated protrusions, we used the fluorescent markers CAAX-HaloTag (plasma membrane) and Actin-HaloTag (actin cytoskeleton). During protrusion formation, the intensity of the plasma membrane probe is increased at the site of protrusion while decreasing elsewhere (Fig. 3A-D, Fig. S6A-F; movie S8). Similar intensity shifts were seen for the actin probe (movie S9). To characterize the flows of membrane and actin over time, we employed Optimal Transport theory (24, 25), where a friction-like dissipation cost and mass displacement are minimized to attain an ‘optimal’ redistribution process (see Supplementary Text; Fig. 3E, Fig. S6G). Our analysis revealed the presence of a cell-wide flow of both plasma membrane and actin cortex toward the protruding front during light-induced protrusion and reversing in direction during recovery (Fig. 3F-I and movie S10, 11). These directed flows provide a potential mechanism to mediate tension propagation following cell protrusion.

**Fig. 3.**
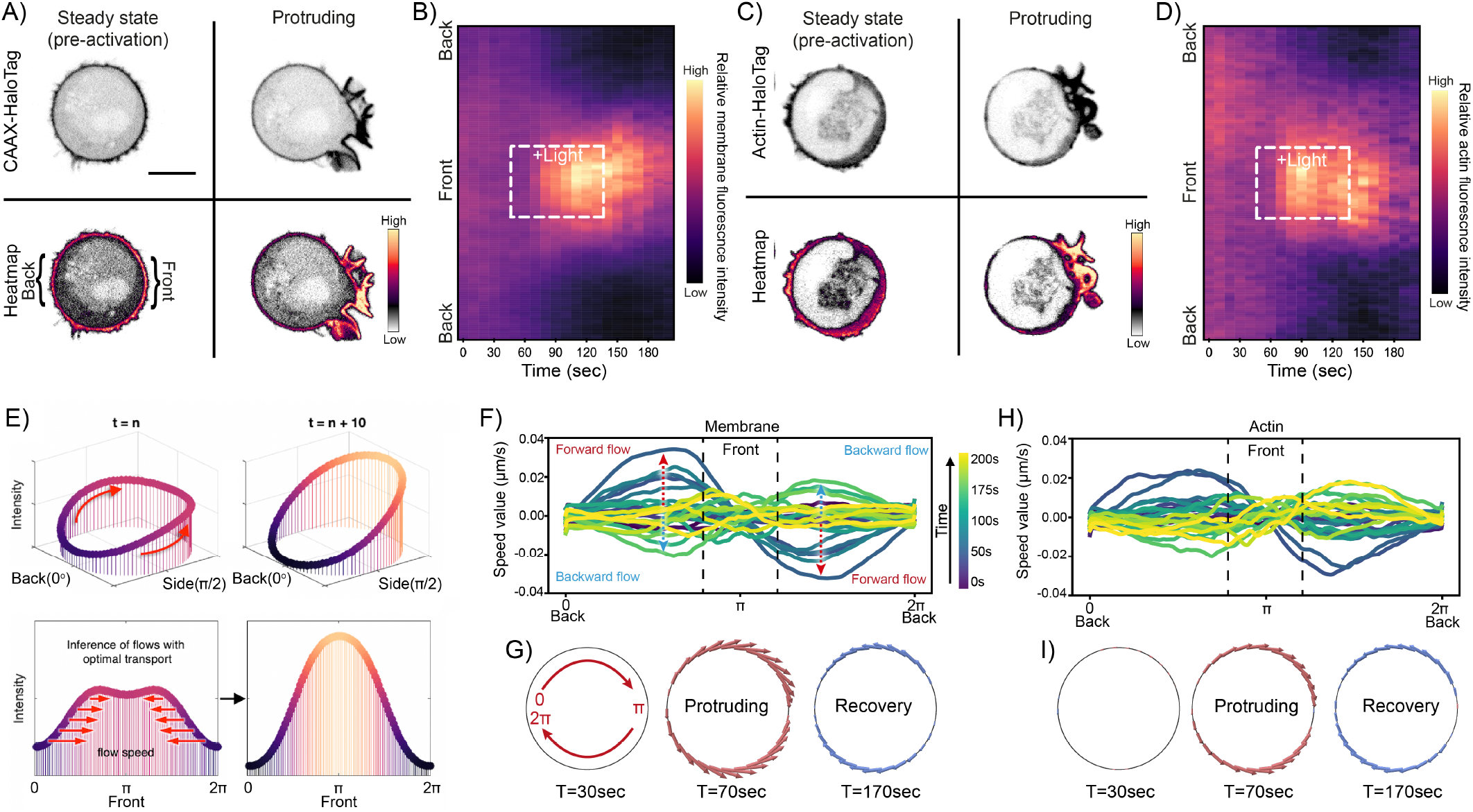
Long-range tension propagation coincides with directed membrane and actin flows toward the protrusion. (A) Confocal images of opto-PI3K cells expressing membrane marker (CAAX-HaloTag): before and during light-activated protrusion. Scale bars: 5 *μ*m. (B) Kymographs of membrane fluorescence along the normalized cell circumference (y-axis) show that over time (x-axis) membrane accumulates towards the protruding cell front and is depleted from the back (n>50, N=6; Fig. S6; see Methods). (C)(D) Similar to (A)(B) but with actin marker (Actin-HaloTag; n>30 N=6). (E) Flows (red arrow) of membrane and actin (intensity plotted in 3D around cell periphery) during protrusion (e.g., t=n to n+10) are calculated assuming optimal transport (see Methods). (F) Membrane flow field inferred from kymograph intensity change over time, assuming optimal transport: shortly after activation begins (t=70s, dark teal traces), the magnitude of membrane flow speed increases (red dashed arrows), with positive speed for clockwise flow along the cell upper half and negative speed for counterclockwise flow along the bottom half (panel G), all moving towards the cell protruding front (*π*). During recovery (t=170s, light green traces), the direction of membrane flow reverses (blue dashed arrows). (G) Membrane flow around the cell before, during, and after (t=30, 70, 170s) right-side protrusion; the flow magnitude is denoted by the arrow size (red: forward flow, blue: backward). Membrane flows toward the protrusion in the protruding phase and away from the protrusion during the recovery phase. (H)(I) Similar to (F)(G) but characterizing actin flow during light-activated cell protrusion.

To infer the critical requirements for cellular membrane tension propagation, we constructed a simple composite mechanical model in which an elastic plasma membrane is coupled to a viscous and contractile gel-like actomyosin cortex (26) via adhesive linkers (Fig. 4A; see Supplementary Text). The membrane displacement (*χ*i) —as a measure of tension propagation upon cortical flows (*υ*i)—is determined by the overall friction imposed through the interconnecting layer of adhesive linkers (e.g., membrane-to-cortex attachment proteins). This friction, *μ*, exerts a drag force on the cell membrane with a magnitude that is proportional to the relative tangential velocity between the cortex and the membrane. Given a moderate membrane-cortex friction, this model adequately captures the known tension responses upon different types of mechanical perturbations (Fig. 4B), including the absence of tension transmission when only the membrane is pulled (e.g., exogenous tether pulling) and rapid propagation upon actin-driven cell protrusion (e.g., endogenous force generation). Furthermore, the model suggests that perturbations engaging both membrane and cortex not only lead to tension propagation but also exhibit a robust tension transmission over a much wider range of membrane-cortex coupling conditions than perturbing either component alone (Fig. 4C). This prediction implies that the key determinant of long-range membrane response is not the endogenous or exogenous application of force but rather whether the mechanical forces directly engage the actin cortex. Putting this prediction to the test, we implemented micropipette aspiration to apply mechanical pulling on both the actin cortex and plasma membrane and monitored tension propagation using our dual-tether assay (Fig. 4D). We detected a rapid, robust, and global increase in membrane tension with little to no attenuation across the cell (Fig. 4E-F, Fig. S7; movie S12-13). Our unifying model indicates that the plasma membrane and actin cortex act as an integrated system for robust membrane tension propagation (Fig. 4G).

**Fig. 4.**
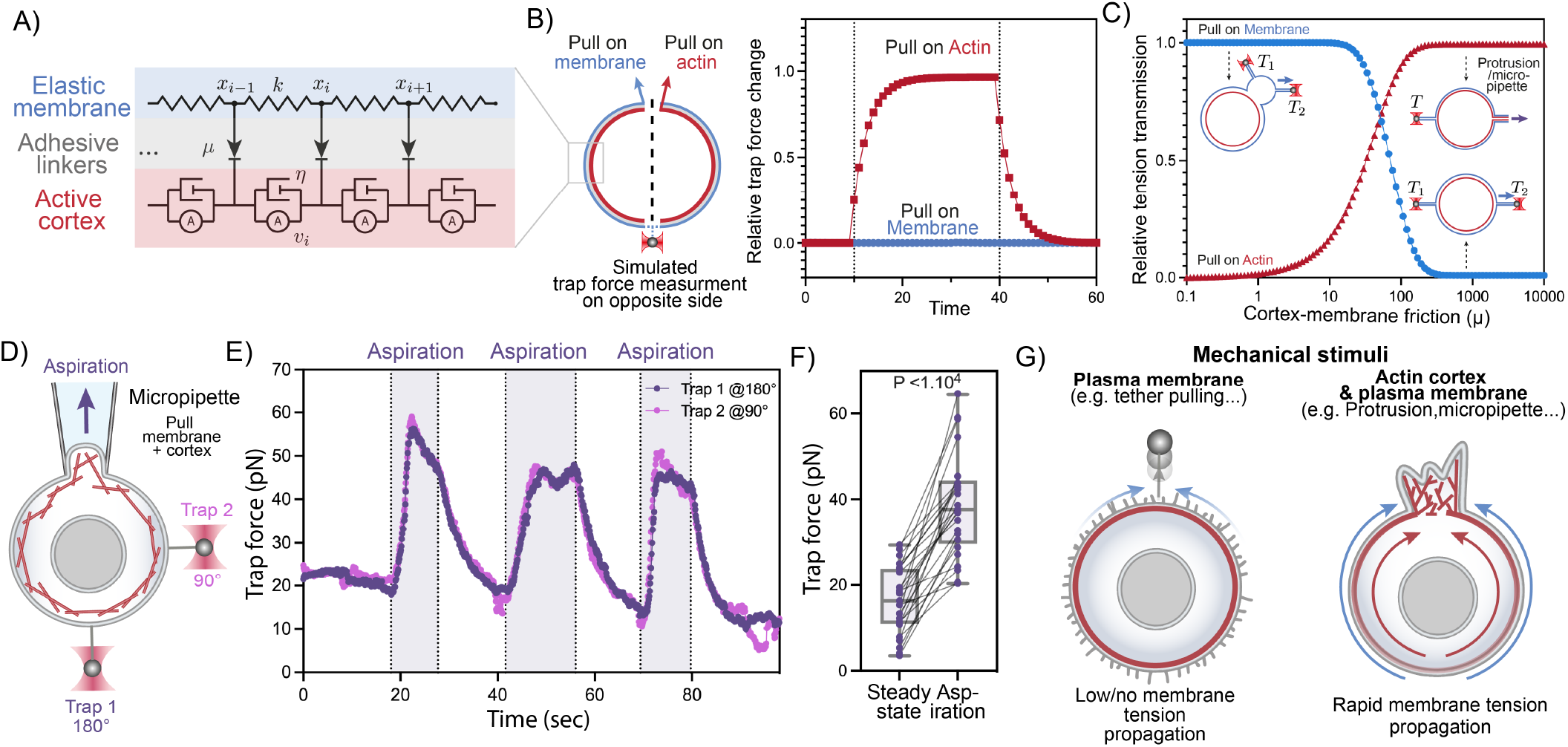
Mechanical forces acting jointly on actin cortex and plasma membrane drive robust rapid long-range tension propagation in cells. (A) A three-tier composite model for membrane tension propagation in cells: membrane displacements (*χ*ii) as a readout for tension propagation upon cortical flows (*υ*i) depend on the membrane elasticity (*κ*) and the membrane-cortex friction *μ* imposed through the adhesive linkers. (B) Model predictions of membrane tension response at moderate membrane-cortex friction (see Supplementary Text): only actin-based pulling leads to tension increase and propagation (red rectangles); external pulling on the membrane alone is inefficient (blue circles). (C) Predicted membrane tension transmission as a function of membrane-cortex friction (x-axis) for different targets of force application: plasma membrane only (blue) and actin cortex only (red). (D) A dual-tether assay to simultaneously monitor membrane tension on the far-end (bottom, trap 1 at 180°) and on the side of the cell (right, trap 2 at 90°) during micropipette aspiration (top), which mechanically pulls on both the membrane and underlying actin cortex. (E) Representative time traces of dual trap forces over successive cycles of aspiration (shaded area) show coinciding tension increases and decreases on both membrane tethers, similar to that in Fig. 2B. (F) Averaged trap forces measured before (Steady-state) and during aspiration. The robust increase in membrane tension upon aspiration on both membrane and cortex is consistent with our model prediction (panel B). Box and whiskers: median and min to max; p values from Wilcoxon paired Student’s t test (n>25, N=5). (G) Schematic of effective membrane tension propagation: in the presence of membrane to cortex attachments, force application to plasma membrane alone does not generate tension propagation, in agreement with the picket fence model. However, mechanical stimuli acting on both actin cortex and plasma membrane lead to rapid and long-range membrane tension propagation across the cell.

## Discussion

By combining optogenetics for local endogenous control of cell protrusion and optical trapping for direct membrane tension measurements in tether pulling assays, we demonstrate that local mechanical force generation such as cellular protrusions elicit rapid long-range propagation of membrane tension throughout the cell. In addition, our findings resolve the long-standing dispute on whether the actin cortex facilitates or impedes tension propagation. When forces are applied to membranes alone (e.g., tether pulling), the actin cortex opposes membrane flow and tension propagation. However, when forces are applied to both the plasma membrane and actin cortex —either during cellular protrusions or upon micropipette aspiration—membrane tension rapidly propagates nearly undampened across the cell. Importantly, our work indicates that membrane tension has the properties expected for a long-range integrator of cell physiology, such as the competition among multiple protrusion sites for a winner-take-all during cell migration (10–12). We speculate that cells may regulate their operational range of membrane tension propagation depending on both the origin of mechanical forces as well as the continuity of the cortex.

## ACKNOWLEDGEMENTS

The authors thank P. J. Zager and M. Wu for kindly sharing plasmids and reagents, and Prof. S. X. Liu for careful reading and comments on our manuscript. We thank all present and past member of the Weiner, Turlier, and Bustamante labs for critical discussions. We also thank the support from Lumicks on C-trap® application in live cell studies, specifically Drs. S. Leachman, N. Hadizadeh, H. Kelkar, and J. Janmaat on the technical assistance in instrumentation, and Drs. E. Lissek, W. Peutz, M. Johnson, and P. Wheeler on the operational sustainability.

Funding: National Institutes of Health grant GM118167 (ODW), National Science Foundation/Biotechnology and Biological Sciences Research Council grant 2019598 (ODW), the National Science Foundation Center for Cellular Construction (DBI-1548297, ODW), and a Novo Nordisk Foundation grant for the Center for Geometrically Engineered Cellular Systems (NNF17OC0028176, ODW), the Labex MemoLife, France under the program “Investissements d’Avenir” ANR-10-LABX-54 (HBR), the European Research Council (ERC) under the European Union’s Horizon 2020 research and innovation programme (Grant agreement No. 949267) (HBR, HT), a QLife (ANR-17-CONV-0005) / QBio grant (ODW, HT), EMBO ALTF 203-2021 (HDB), National Institutes of Health grant K99GM137074 (SY), and Howard Hughes Medical Institute (CB).

## Materials and Methods

### Cell culture & cell lines

HL-60s were cultured in R10 growth medium, which is RPMI 1640 supplemented with L-glutamine and 25 mM HEPES (Corning; Corning, NY) and containing 10% (v/v) heat-inactivated fetal bovine serum (Gibco; Waltham, MA). Cultures were kept at a density of 0.2–1.0 million cells/mL at 37°C/5% CO_2._ HEK293T cells (used to make lentivirus for transduction of HL-60s) were grown in DMEM (Corning; Corning, NY) containing 10% (v/v) heat-inactivated fetal bovine serum (Gibco; Waltham, MA) and maintained at 37°C/5% CO_2_. All media were 0.22-um filtered.

Opto-PI3K cells (iLid-BFP-CAAX, iSH2-GFP, Pak-PBD-mCherry) were obtained from (*1*). Opto-PI3K cells expressing either the membrane marker CAAX-HaloTag or the actin marker Actin-HaloTag were generated using constructs kindly provided to us by Patrick Zager.

### Transduction of HL-60 cells

HEK293T cells were used to generate lentivirus and were seeded into 6-well plates until grown at approximately 80% confluent. For each transduction, 1.5 μg pHR vector (containing the appropriate transgene), 0.167 μg vesicular stomatitis virus-G vector, and 1.2 μg cytomegalovirus 8.91 vector were prepared for transfection using TransIT-293 Transfection Reagent (Mirus Bio; Madison, WI). Three days post transduction virus-containing supernatants were harvested and concentrated approximately 40-fold using Lenti-X Concentrator (Clontech; Mountainview, CA). Concentrated viruses were frozen and stored at −80°C until needed. For all transductions, thawed virus was mixed with approximately 0.3 million cells in growth media supplemented with polybrene (8 μg/mL) and incubated overnight. Cells expressing desired transgenes were isolated using fluorescence-activated cell sorting (FACS) as appropriate (FACSAria2; BD Biosciences; Franklin Lakes, NJ).

### Microscopy hardware

Imaging depicted in Fig. 1B, 3B, 3C, S6A, S6C and Movie S1, S6, S7 were performed at 37°C on a Nikon Eclipse Ti inverted microscope equipped with a Borealis beam conditioning unit (Andor), a CSU-W1 Yokogawa spinning disk (Andor; Belfast, Northern Ireland), a 100X PlanApo TIRF 1.49 numerical aperture (NA) objective (Nikon; Toyko, Japan), an iXon Ultra EMCCD camera (Andor), and a laser merge module (LMM5, Spectral Applied Research; Exton, PA) equipped with 405, 440, 488, and 561-nm laser lines. All hardware was controlled using Micro-Manager (UCSF).

### Preparation of Opto-PI3K cells for confocal imaging

For experiments in which we monitored cells by confocal imaging, cells were seeded in a 96-well #1.5 glass-bottom plates (Brooks Life Sciences; Chelmsford, MA) in R+B imaging media, which is RPMI 1640 supplemented with L-glutamine and 25 mM HEPES (Corning; Corning, NY) and containing 0.2% Bovine Serum Albumin (BSA, endotoxin-free, fatty acid free; A8806, Sigma; St. Louis, MO). For plasma membrane and actin imaging using HaloTag (Fig 1B, 3A, 3C), cells were stained with 100nM of JF646X for 10 min before being pelleted at 300*g* for 3 min and resuspended in R+B imaging media (RPMI+0.2% BSA). For plasma membrane imaging using the membrane dye CellMask, cells were first incubated with ∼2-5μg/ml of CellMask™ Deep Red (C10046, Thermofisher) for 3 minutes at 37°C/5% CO_2_. Cells were then pelleted at 300*g* for 3 min and resuspended in R+B imaging media (RPMI+0.2% BSA).

### Preparation, settings, and operation procedures for membrane tethering pulling experiments on C-trap^®^ optical tweezers with confocal imaging

#### Cell preparation

Opto-PI3K: 1-1.5 ml cells (from culture at density of 0.6–0.8 million cells/mL) were stained (with 0.5 μl of CellMask™ Deep Red or 100nM of JF646X) in the absence of presence of actin inhibitor: 10 μM Latrunculin B; then pelleted down and resuspended in either R+B imaging medium (RPMI+0.2% BSA) or R10 medium (all media 0.22-um filtered) for samples used in tether pulling assay.

#### Bead preparation

In a 1.7 ml Eppendorf tube, the following solutions were added: 9 μl of ultrapure water (Corning, 46-000-CM), 9 μl of carboxyl latex bead (4% w/v, 2 *μ*m; Invitrogen, C37278), and 2 μl of Concanavalin A (1 mg/ml; Sigma-Aldrich, C2272); sample was vortexed at low speed at room temperature for 45-60 min; 1-2 μl of this bead mixture stock was added into 1 ml of RPMI 1640 buffer (0.22-um filtered) for samples used in tether pulling assay.

#### Microfludics

An u-Flux™ flow cell (70-mm chips; Lumicks, C1), installed on an heat-insulating PVC holder, was passivated with R+B imaging media (0.22-um filtered) and pre-warmed at 35-37°C for 1-2 hours. A custom-made microchamber integrated with micropipettes (descriptions on the assembly provided at the end) was used in place of u-Flux™ flow cell to apply aspiration in tether pulling assay performed on C-trap^®^ (instrument operation procedures in the next section). During the assay, an air-pressured microfluidics flow system (u-Flux™, Lumicks), with pre-cleaned and proper dimensions of tubing connections, was used to deliver cell samples, bead solutions, and blank media/buffers (for flushing) into the flow cell or microchamber. Specifically, a tubing with large ID (1/32 inch; Idex, 1520L) was used to deliver cells at the lowest pressure setting (0.04-0.12 mbar, or sometimes just gravity flow) so as to minimize the shear force exerted to the cells during delivery. The delivery of beads and media was made with a narrower tubing (ID 0.004 inch; Upchurch Scientific, PM-1148-F). After flowing ∼200-500 μl (sufficient to displace dead volumes combined within the microfluidics system) of cells into the C-trap^®^ system pre-warmed at ∼36°C, incubated for 15-20 min so that the cells settle and stably attach to the bottom surface of the u-Flux™ flow cell, then cell locations were marked prior to the subsequent tether pulling experiments with optical traps. The cell samples were replenished every 1.5-2 hours, with abundant flushing of R+B imaging medium in between (which ensures the flow cell surface remains properly passivated).

#### Optical trapping – setting and operations

A commercial dual-trap optical tweezers with 3-color confocal imaging, aka C-trap^®^, from Lumicks was used to perform the tether pulling assay with concurrent fluorescence imaging. The flow cell, or microchamber, held paralleled to the table surface was aligned perpendicular between a water objective (60x, NA 1.2; Nikon, MRD07602) coming from the bottom and a matching condenser (60x, NA 1.4, used with Type A immersion oil; Leica) coming from the top. The flow cell was positioned in between the two such the IR laser beams (1064 nm) focused down by the objective were formed inside the flow cell ∼10-20 μm above the inner bottom surface (with the flow cell nano-stage set at the middle position), so the condenser can adequately collect photons from the IR trapping beams post flow cell and project on to position-sensitive detectors (PSDs) for accurate trap force measurement (data acquisition at a rate of 78125 Hz and later down sampled to 10 Hz for analysis). The objective also directs fluorescence excitations in the visible wavelength range (488, 532, and 642 nm respectively for opto-tool, Rac biosensor, and CellMask/HaloTag) into the flow cell (or microchamber). The two set of light sources (IR and visible) were controlled by separate telescopes and mirror-steering systems upstream from the objective. The same objective collected the emission photons from the imaging/optical trapping sample plane inside the flow cell for fluorescence imaging (bandpass filters: 512/25, 582/75, and 680/42; camera pixel size: 100 nm; frame rate depends on confocal scanning area size), whereas the condenser provided bright field imaging (850 nm LED light source) recorded at 10 Hz.

Both the objective and the condenser were pre-warmed to 35-37°C (temperature control unit, Lumicks) for at least 2-3 hours prior to cell experiments. The IR trapping power was typically set at 100% trapping laser, 10% overall power, and 50-50 split between trap 1 (T1) and trap 2 (T2), which is about ∼175 mW per trap (measured at the objective front) and ∼0.2 pN/nm in trap stiffness for a 2-μm bead (bead corner frequency ∼2500 Hz). Low settings of excitation laser were sufficient for fluorescence imaging (typically ∼2-5% of total power gives ∼0.02-0.04 μW measured at the objective front), minimizing the photo-toxicity to the cell during experiments.

At the beginning of each cell recording in the tether pulling assay, 2-μm Concanavalin-coated beads were perfused into the assay (e.g., at 0.4 mbar via channel 5 in u-Flux™ C1 flow cell) and single beads were captured in either one or both traps, and the flow cell stage was moved to bring the beads to a location previously pre-marked. With beads in the vicinity of the cell, i.e., in z-axis at the same confocal imaging plane for the cell (∼2-6 μm from the flow cell bottom surface) and ∼4-6 μm away from the cell body in the x-y plane, the trap stiffness was calibrated and any residual force readout were zeroed before engaging the bead with the cell body to form membrane tethers. Region of interest (ROI) was cropped for bright field imaging (typically an area of 35×45 μm) continuous recording at 10 Hz was initiated.

1. Tether pulling assay with light-activated cell protrusions: as seen from the bright field camera, we approached beads to position them in direct contact with the cell body (even pressing a little, judging from the counter force acted on the bead in the trap), we waited for several seconds, and then we carefully pulled out membrane tethers (∼4-10 μm in length) at the desired configuration (e.g., two tethers right angle from each other). We monitored steady state tension for at least 1 min (Fig. S2A-C) before illuminating with local 488-nm excitation (ROI: 6×10 μm) continuously for 90 sec on the opposite site of (or right angle from) the membrane tether. Upon localized 488-nm illumination, the local recruitment of opto-controls (iSH2 labelled with EGFP) to trigger cell protrusions was also imaged simultaneously (∼1-1.3 sec/frame scanned). Post protrusion activation, we monitored cell membrane tension recovery for 180 sec and repeated activation cycles for as long as the tethers last (see Movie S4-5). At desired time points, i.e., before, during, or after 488-nm light activation, the activated Rac was specifically imaged via 532-nm illumination to visualize the distribution of the Rac biosensor (Pak-PDB-mcherry) inside the cell (see Fig. S1C-E; Movie S3). Similarly, the changes in cell membrane morphology were imaged over time with 642-nm illumination (for CellMask Deep Red or Halo-tag 660 if cells were stained earlier).
2. Other experimental conditions in tether pulling assay, including controls: following the same bead engagement procedure described above, membrane tethers were pulled out from the cell body or from small patches of vesicle-like, outward budding membrane blebs that are detached from actin cortex upon Latrunculin treatment. Specifically, after the first membrane tether was pulled, the second tether was pulled from a nearby location ∼2 μm away. The membrane tension was recorded in the same fashion as detailed earlier but for the following conditions: light activation on wild-type cells or drug-treated opto-PI3K cells; in the absence of any light illumination, we moved one trap to extend the length of one tether on the cell body (or bleb) and monitor the tension response on the other (see Movie S6-7); or instead of 488-nm illumination (which triggers actin-driven cell protrusion) the cell was engaged with micropipette aspiration, which exerts mechanical pulling on both membrane and cortex, and the membrane tension was recorded over cycles of aspiration and relaxation (see below; Movie S12-13).

#### Micropipette aspiration

A custom-made microchamber was used to implement micropipette aspiration on the C-trap^®^ system. Specifically, a micropipette of 2-6 um tip diameter was prepared by gravity pulling a thin glass capillary tube (ID 0.040 +/- 0.010 mm, OD 0.080 +/- 0.010 mm, length 150 mm; King Precision Glass, KG-33) that was threaded through a heated platinum coil (∼2 mm in diam.; Pt wire is 0.3 mm in diam., Alfa Aesar) upon application of a desired voltage. The micropipette tip size generally correlates with the heating time required to pull the glass tube apart; the faster heating, the more rapid the pull, giving micropipette tips in smaller diameters. The micropipette was then sandwiched between a 1-mm glass cover slide (3”x1”x 1mm; VWR) and a #1.5 glass cover slip (24×60 mm; VWR), held together with two pieces of melted Nescofilm (100 um in thickness; Karlan) as sealant and spacer in the microchamber. 6 holes were drilled prior on the glass slide to provide inlets, which are connected to valves and uFlex™ pressurized syringe reservoirs (for cell and bead samples delivery as well as buffer flushing), and outlets towards the waste collection. The micropipette was connected to a separated microfluidic pressurized system (MFCS™-EZ from Fluigent; input: -600 mbar, output: -69 to 0 mbar) powered by a small floor pump (KNF, model: N86KN.18, with manual regulator) to provide aspiration control in the tether pulling assay on C-trap^®^. The aspiration pressure zero point for each micropipette was carefully calibrated and set to have no outward nor inward flow detectable to a laser trapped bead that was placed at the tip opening of the micropipette. During the experiments, cells were delivered into the microchamber at the same gentle flow rate (0.04-0.12 mbar, or sometimes just gravity flow) and captured by the optical trap, which quickly brings the cell to the micropipette tip. A minute amount of suction was applied to keep the cell stably engage with the tip (so it neither floats away from the tip nor falls back into the optical trap) but without any visible deformation of the cell morphology (as seen in bright field camera). Then following the same bead calibration procedure and membrane tether pulling process as described earlier, consecutive rounds of aspiration and relaxation were performed on the cell for as long as the membrane tethers persist (see Movie S12-13).

##### Image and membrane tension analysis

Fiji (NIH), Excel (Microsoft; Redmond, WA), custom Python code, and Prism (Graphpad software, Inc) were used for image and membrane tension analysis.

Average trap force plots (Fig. 1E, 2C, S3A, B, S4A, B) were obtained by aligning trap force traces at time of light induction.

Average linked trap force plots were made using Prism Graphpad software, Inc). In Fig. 1F, average trace trap force was measured for 60 seconds before light induction (steady state) and for the duration of the light induction (90 seconds, Light). For Fig. 4F, average trace trap force shown here for 30 seconds before aspiration (steady state) and for the duration of aspiration (15-30 seconds) and intervening recovery periods.

Pearson correlation coefficients between T1 and T2 were calculated using Prism Graphpad software, Inc) (Fig. 2H, Fig. S4D, Fig. S5J, and S8G). For Fig. 2H and S5J, we used 30 seconds before activation for steady state, 30 seconds of light induction for opto-activated protrusion, and ∼10-30 seconds of active tether pulling on cell membrane (tether length >30 μm) and on blebs. In Fig. S4D, for ‘+Light’ we used the full duration of light activation (90sec) and for ‘Recovery’ 70sec post light induction. In S8G we used 15-30 sec pre-aspiration for steady state value and full duration of aspiration (∼15-30sec) for aspiration.

Delay between T2 and T1 during light induced protrusion (Fig. S4E) was calculated by measuring the time difference between light induction and change in trap force slope for each trap. Of note, measuring time difference from light induction to plateau in force increase yields similar results (i.e., delay time between the two traps is still of ∼ 1 sec).

For measurement of relative tether force over distance of moving tether (Fig. S5K), we normalized the trap force of static tether by its average when the extending tether was at distance <1 μm (namely, before any active pulling).

##### Statistical analysis

For all statistical analysis, PRISM 9 (Graphpad software, Inc) was used. Statistical details can be found in the legend of each figure. N represents number of independent biological replicates. Pooled independent experiments are used in dot plots.

## Supplementary Text

**Movie S1.**

Time-lapse confocal images of HL-60 cells expressing opto-construct (Opto-PI3K) and membrane marker (CAAX-HaloTag) showing localized membrane protrusion upon light activation. Related to Fig. 1B. Scale bar: 5μm.

**Movie S2.**

Time-lapse (confocal) of Opto-PI3K HL-60 cell expressing Rac biosensor Pak-PBD-mCherry undergoing light-induced protrusions, confirming active Rac recruitment at the site of light-induction. Related to Fig. S1B. Scale bar: 5μm.

**Movie S3.**

Time-lapse (confocal, taken on C-Trap) of Opto-PI3K HL-60 cell expressing iSH2-GFP. Heatmap represents fluorescence signal intensity showing iSH2 recruitment from the cytoplasm to the plasma membrane shortly after light induction. Related to Fig. S1C-E.

**Movie S4.**

Brightfield imaging and time trace of trap force measurement of a protruding cell exhibiting sharp increase in membrane tension during light-activated protrusion on the opposite end of the cell Related to Fig. 1D-E and S3A-C. Scale bar: 5μm.

**Movie S5.**

Brightfield imaging and time trace of dual trap force measurements of a protruding cell showing nearly coinciding tension increases on both membrane tethers adjacent to (trap 2) and at the opposite cell surface from (trap 1) protrusion. Related to Fig. 2A-C and S4A-C. Scale bar: 5μm.

**Movie S6.**

Brightfield imaging and time trace of measurements of a dual-tether assay to detect tension propagation between two tethers closely pulled out from the cell body membrane. The bead on the right continuously extends the tether on the right, thereby applying mechanical force to the cell membrane. Trap force (as tension responses readout) from the static tether remain uncorrelated to that of the moving tether Related to Fig. 2D-E and S5A-B. Scale bar: 5μm.

**Movie S7.**

Brightfield imaging and time trace of measurements of a dual-tether assay performed on a bleb to detect tension propagation between two tethers closely pulled out from the bleb membrane. The bead on the right continuously extends the tether on the right, thereby applying mechanical force to the bleb membrane. Related to Fig. 2F-G and S5C-D. Scale bar: 5μm.

**Movie S8.**

Time-lapse (confocal) of Opto-PI3K cells expressing membrane marker (CAAX-HaloTag) undergoing light-induced protrusions, related to Fig. 3B. Scale bar: 5μm.

**Movie S9.**

Time-lapse (confocal) of Opto-PI3K cells expressing actin marker (Actin-HaloTag) undergoing light-induced protrusions, related to Fig. 3D. Scale bar: 5μm.

**Movie S10.**

Membrane flow around the cell before, during, and after right-side protrusion; the flow magnitude is denoted by the arrow size (red: forward flow, blue: backward). Membrane flows toward the protrusion in the protruding phase and away from the protrusion at the recovery phase. Related to Fig. 3G.

**Movie S11.**

Actin flow around the cell before, during, and after right-side protrusion; the flow magnitude is denoted by the arrow size (red: forward flow, blue: backward). Actin flows toward the protrusion in the protruding phase and away from the protrusion at the recovery phase. Related to Fig. 3I.

**Movie S12.**

Brightfield imaging and time trace of trap force measurements during micropipette aspiration, which mechanically pulls on both the membrane and underlying actin cortex. Scale bar: 5μm.

**Movie S13.**

Brightfield imaging and time trace of dual trap force measurements during micropipette aspiration, which mechanically pulls on both the membrane and underlying actin cortex. Scale bar: 5μm.

**Fig. S1.**
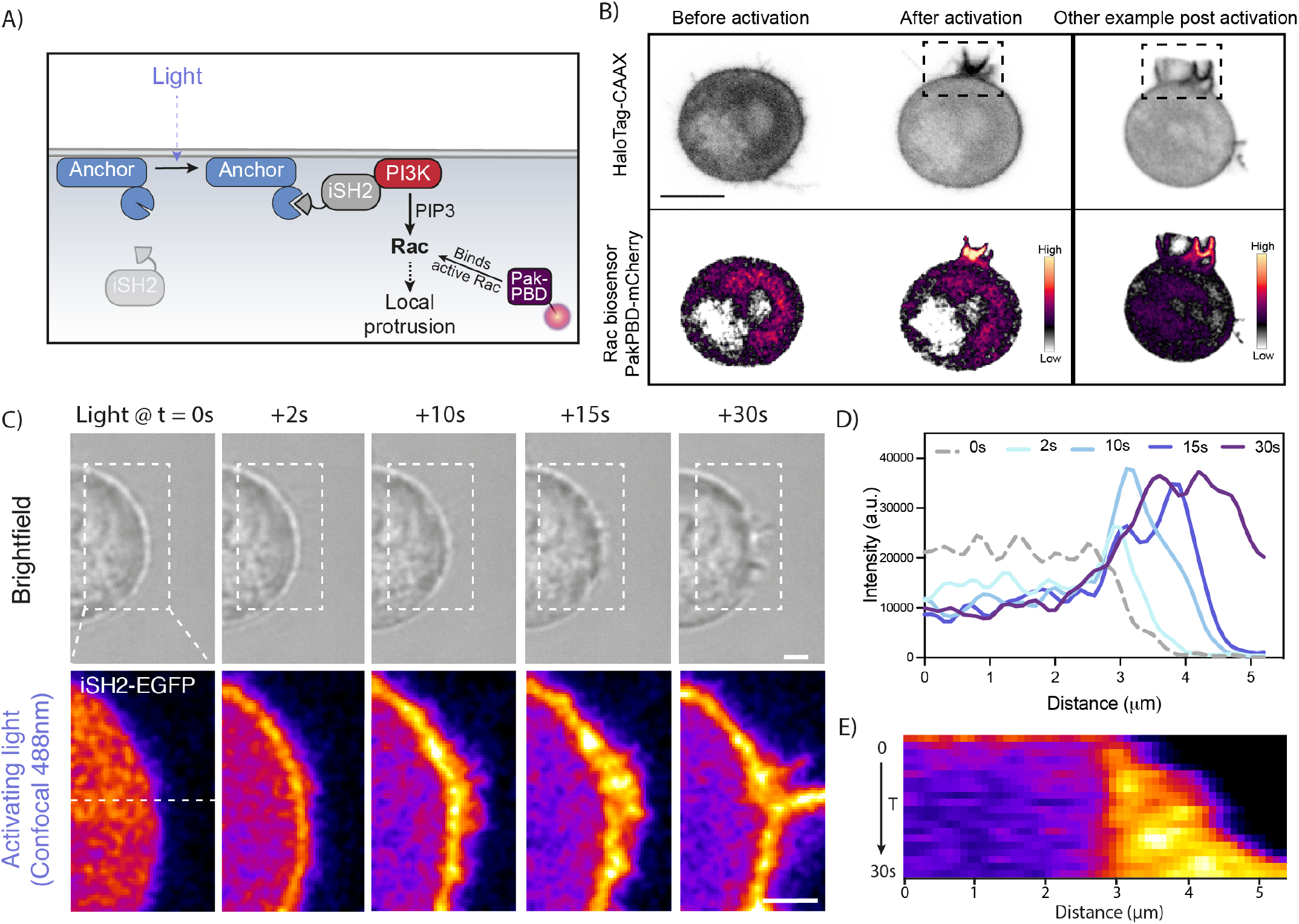
Optogenetic control of PI3K leads to local Rac activation and triggers localized actin-driven cell protrusion. (**A**) Membrane-anchored optogenetic control for light-induced activation of phosphoinositide 3-kinase (PI3K): upon localized 488-nm excitation, the membrane anchor protein (iLid-BFP-CAAX) undergoes a conformational resulting in the binding and recruit inter SH2 domain (iSH2) to the illuminated region. iSH2 proceeds to recruit PI3K, whose lipid product (PIP_3_) induces the activation of Rac GTPase (Rac). Active Rac then triggers actin polymerization leading to localized membrane protrusion. By imaging the mCherry-labelled Rac biosensor (Pak-PBD-mCherry), which recognizes and binds the active GTP-bound Rac, we can monitor Rac activation during light-induced protrusions (see Methods). (**B**) Time-lapse confocal images of HL-60 cells expressing opto-construct (Opto-PI3K), membrane marker (CAAX-HaloTag, imaged on top), and Rac biosensor (PAK-PBD-mCherry, imaged on bottom). Middle and right: localized recruitments of active Rac is confirmed at the site of light activation for cell protrusion (box in black dashed line). (**C**) Time-lapse brightfield (top) and confocal images (bottom) of an opto-PI3K cell during light activation. The specific recruitment of PI3K activator, (iSH2-EGFP) to the illuminated area (box in white dashed line) is monitored upon 488-nm excitation. Within 2 s (between the first two frames), iSH2 has redistributed from the cytoplasm to the plasma membrane. Scale bar: 1μm. **(D)** Fluorescence intensity line scans (along the white dashed line in panel C) show the enrichment of opto-construct (iSH2-GFP) at the cell protruding site over time. (**E**) Kymograph of the above line scan (white dashed line in panel C) shows that after iSH2 is recruited to the membrane, the cell contour (i.e., its membrane) rapidly expands outward.

**Fig. S2.**
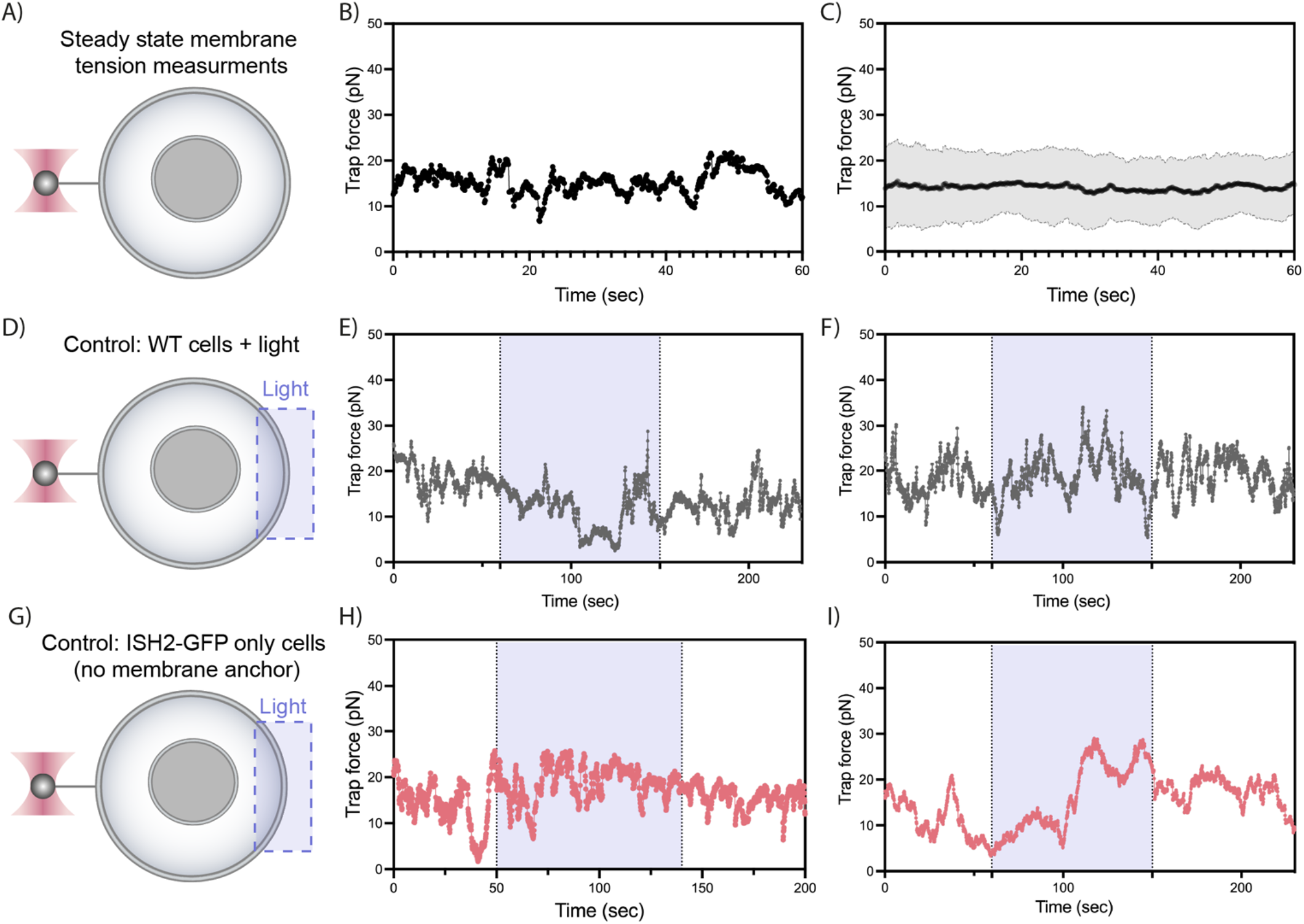
Membrane tension measurements controls. (**A**) An optical trap tether pulling assay to monitor basal membrane tensions without any light-activated cell protrusion, thus termed tension at “Steady state.” The magnitude of cell membrane tension is probed via a membrane tether between the cell body and a 2-um latex bead held by an optical trap (see Methods). Cell membrane under tension exerts a force to the trapped bead through the membrane tether, and this force causes a displacement of the bead center relative to the optical trap center, providing a direct readout of membrane tension. **(B)** Representative time trace of trap force measured from the tether pulling assay with a cell at steady state: membrane tension remains stable with low magnitude of stochastic fluctuations. **(C)** Averaged time trace of trap force for cell membrane tension recorded at steady state (means ± SDs; n>50, N=8). **(D)** As a control, we light activate the wild-type (WT) cells, which lack opto-constructs, and use the same tether pulling assay described above to monitor membrane tension response before, during, and after 488-nm illumination (purple shaded area). **(E)(F)** Representative time traces of trap force for cell membrane tension recorded from WT cells with light activation. The activation light alone does not elicit any changes in cell morphology or membrane tension responses. **(G)** In another control, we light activate cells lacking the membrane anchor protein for opto-control (iLid-BFP-CAAX) and monitor their membrane tension response upon 488-nm illumination (purple shaded area). **(H)(I)** Representative time traces of trap force for cell membrane tension recorded from cells with lacking expression of one or more opto-constructs. Same as panel E-F, no perceptible changes in cell morphology or membrane tension were observed.

**Fig. S3.**
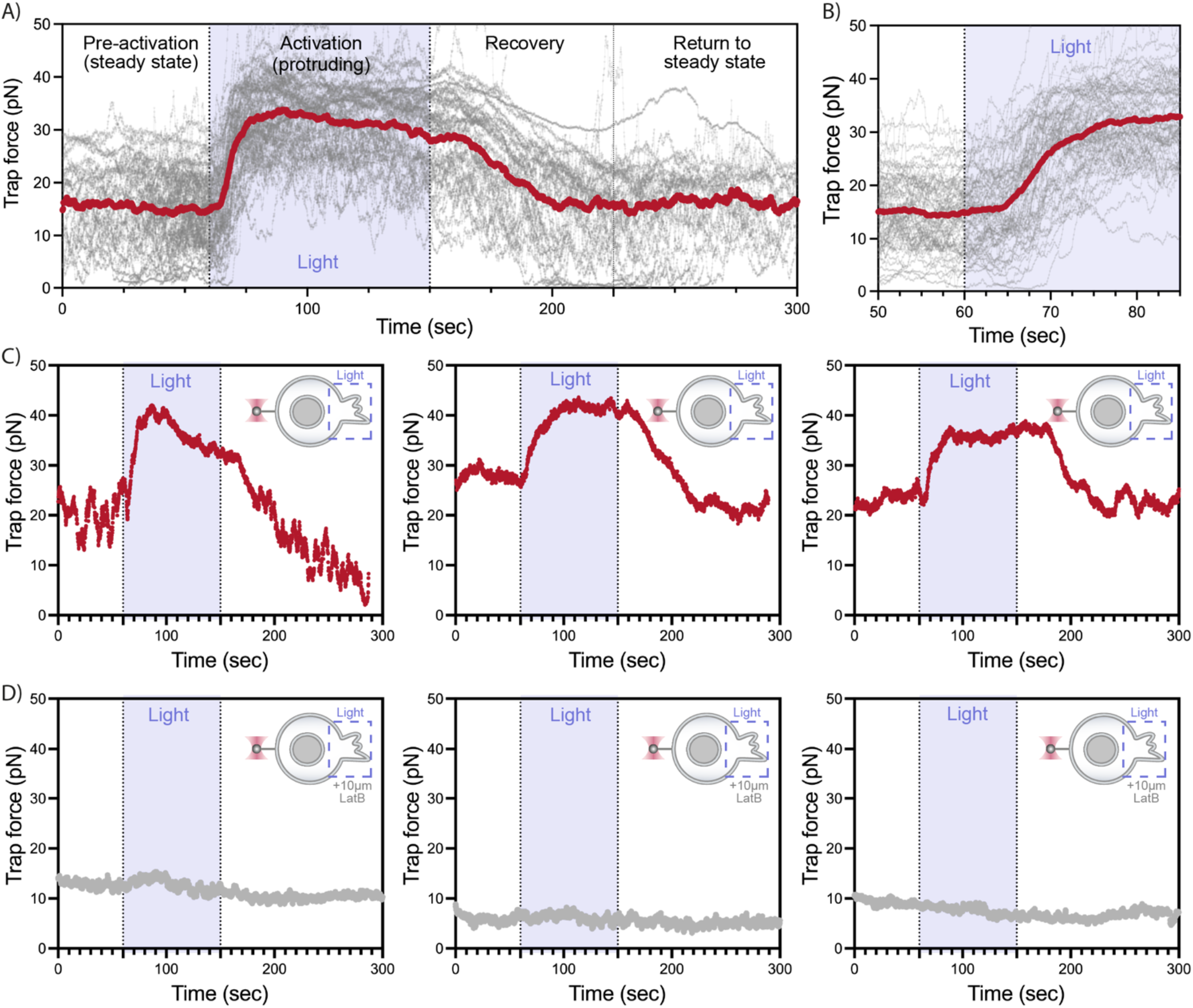
Membrane tension robustly increases at the opposite site of the cell during actin-driven protrusion. **(A)** Averaged time trace of trap force (red) for cell membrane tension recorded before (steady state), during (activation), and after (recovery and return to steady state) light-induced protrusion on the opposite side of the cell (see Fig. 1C). Individual data traces are shown in light grey (n>60, N=8). Cells at steady state exhibit stochastic fluctuations in membrane tension, similar to that shown earlier in Fig. S2B-C. Upon light activation (purple shaded area), membrane tension rapidly increases and levels off to a plateau towards the end of activation (total 90s). The presence of a plateau potentially indicate that membrane reservoirs unfold to provide extra membrane, thus buffering the tension rise. Shortly after the activation light is turned off, membrane tension gradually decreases to the steady state level. **(B)** Zoom-in on traces in panel A: increases in membrane tension emerge within the first 5-10s of light activation. **(C)** Three example time traces of trap force for membrane tension before, during, and after light-induced cell protrusion. **(D)** Same as panel C but recorded from cells treated with actin inhibitor (10μM Latrunculin B). We verified that Latrunculin B treatment neither impairs the opto-tool recruitment nor the subsequent Rac activation. This control shows that the increase in membrane tension measured at the opposite side of cell protrusion (panel A-D) is actin-driven.

**Fig. S4.**
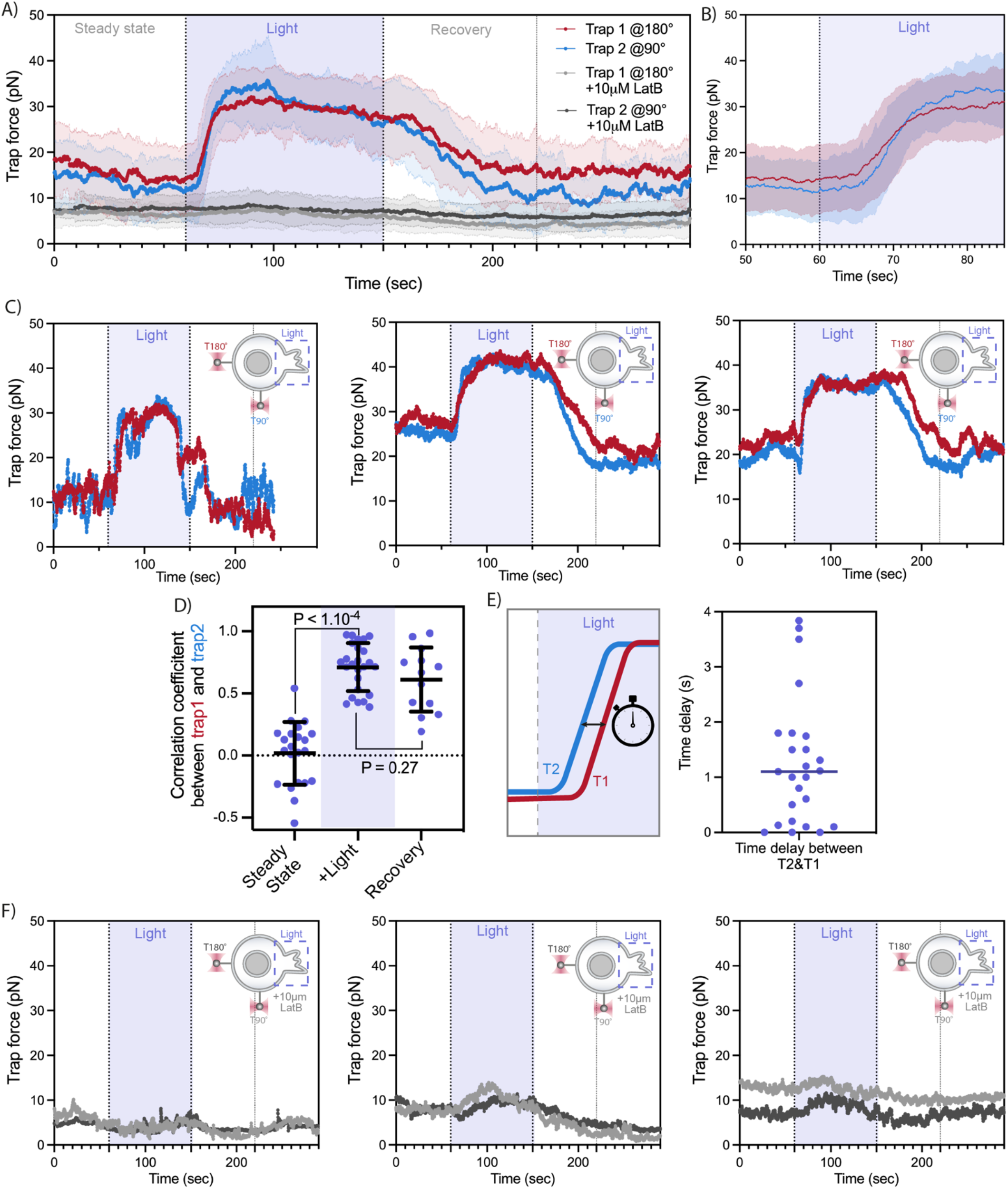
Membrane tension propagates within seconds across the cell during actin-driven protrusion. **(A)** Red and blue: averaged time traces of trap force for dual membrane tension measurements before (steady state), during (Light), and after (recovery) activating cell protrusion. A nearly coinciding tension increase is observed between the membrane tether adjacent to (trap 2, blue) and at the opposite cell pole from (trap 1, red) a protrusion. Grey: as a control, averaged trace from cells treated with actin inhibitor (10 μM Latrunculin B) shows no membrane tension change upon activation (means±SD; n>15, N>4). **(B)** Zoom-in on traces in panel A: increases in membrane tension emerge on both tethers within the first 5-10s of light activation. **(C)** Three example time traces of trap force for dual membrane tension measurements before, during, and after light-induced cell protrusion. At steady state tensions from the two tethers show little correlation, but they become highly correlated upon light activation (purple shaded area). During the recovery phase, we often observe a lag in time between the two tethers’ tension drop, with the tether at the opposite cell surface from the protrusion site recovering slower (red). **(D)** Pearson correlation coefficient between dual trap forces measured at steady state, during light activation, and recovery afterwards (70s post light). Error bar: means ± SD; p values from Welch’s unpaired Student’s t test (n>10, N>4). **(E)** Left: time delay measured between tension rise on membrane tethers adjacent to (trap 2 at 90°, blue) and at the opposite cell pole from (trap 1 at 180°, red) the protrusion. Right: in most cells, the traps detects membrane tension increase on both tethers within a second or less of each other, indicating a rapid propagation of tension across the cell. **(F)** Three example time traces of trap force for dual membrane tension measurements with cells treated with actin inhibitor (10μM Latrunculin B) before, during (purple shaded area), and after light activation of cell protrusion.

**Fig. S5.**
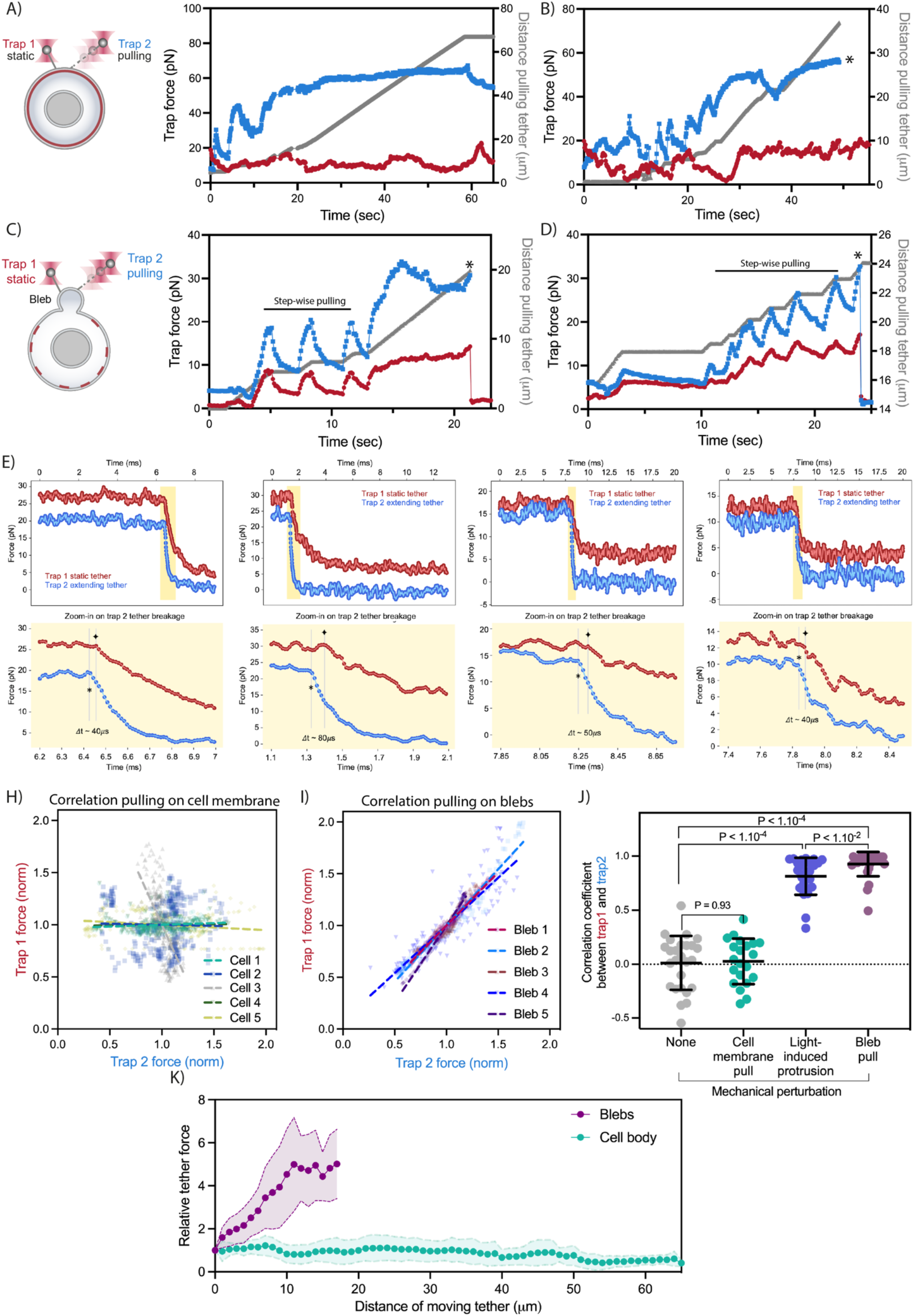
Mechanical perturbations affecting only the plasma membrane do not result in measurable membrane tension propagation in cells but do in blebs detached from actin cortex. **(A)** An example time trace of trap force for dual membrane tension measurements, where one moving trap (T2, blue) mechanically perturbs on the cell membrane by continuously pulling and extending the membrane tether, and the other trap remains static (T1, red) to monitor changes and propagation in tension to a nearby membrane tether. The increase in length of the extending tether from the cell body is plotted in grey along the right y-axis. **(B)** Another example recorded under similar operations as panel A; ‘*’ annotates when the extending tether broke. Note that a sudden tension release upon breakage of the extending tether (blue, at t ∼50s) does not lead to changes in tension on the static tether (red), which is in close proximity to the extending tether (≤ 2μm). This observation shows that mechanical perturbations affecting only the plasma membrane in cells are locally constrained and inadequate to generate measurable tension propagation between the two tethers. **(C)(D)** Similar operations as panel A-B but monitoring tension propagation between two membrane tethers on cellular blebs (i.e., a vesicle-like, small section of membrane detached from actin cortex upon Latrunculin B treatment). The tension readouts between the extending and the static tethers on blebs appear highly correlated, unlike those on cell body in panel A-B. Specifically, during the “step-wise pulling” (jerky motion) to extend tether in trap 2 (blue), the static tether held in trap 1 (red) exhibits immediate spiky rises in tension, mirroring the pattern in trap 2. When a “continues pulling” (smooth motion) is exerted on the extending tether by trap 2 (blue, at t∼18s in panel C), the tension increase on static tether (red) accordingly becomes gradual. Furthermore, the sudden drop in tension back to initial level on the static tether (red, t∼26s)—in response to the sudden tether breakage (*) and thus tension release of the extending tether (blue)—reflects a direct tension transmission and rapid propagation (see panel E-F) within a membrane bleb detached from the constraining actin cortex. **(E)(F)** Example zoom-in traces of dual trap forces (raw data at 78 kHz) showing the time difference between a sudden tension release upon breakage (*) of the extending tether (blue) and the subsequent reduction (✦) in tension on the static tether (red; traces slightly offset in y-axis for illustration clarity). Typically, this time delay observed is ≤ 100 μs (measured between the inflection points, * and ✦, on each trace), which is right around the temporal resolution of our optical trapping instrument (limited by the corner frequency of a 2-μm bead held by a trap with stiffness of ∼0.2 pN/nm), indicating that the actual time scale of tension propagation on cellular blebs is likely too fast to be resolved in our experiments. **(H)(I)** Correlation plots of normalized trap forces between the moving and static tethers: low for membrane tether pulling on cell body (panel H), and high for pulling on vesicle-like cellular blebs (panel I). Five representative measurements from different cells are shown per conditions; dashed lines: linear regression. **(J)** Pearson correlation coefficient between dual trap forces measured before any perturbations (None), during cell body membrane pulling, upon light-activated actin-driven protrusions, and during membrane bleb pulling (some data already plotted in Fig. 2H). Error bar: means ± SD; p values from Welch’s unpaired Student’s t test (n>20, N>3). **(K)** Relative force changes (y-axis) for membrane tension monitored on the static tether as a function of the extending tether length (x-axis) upon continuous pulling. In the case of blebs (purple), the tension on static tether increases as the extending tether lengthens; however, there are no perceptible changes in static tether tension even when the other tether has extended more than 60 μm from the cell body (green) (n>15, N>3). Graphical data represent means ± SDs.

**Fig. S6.**
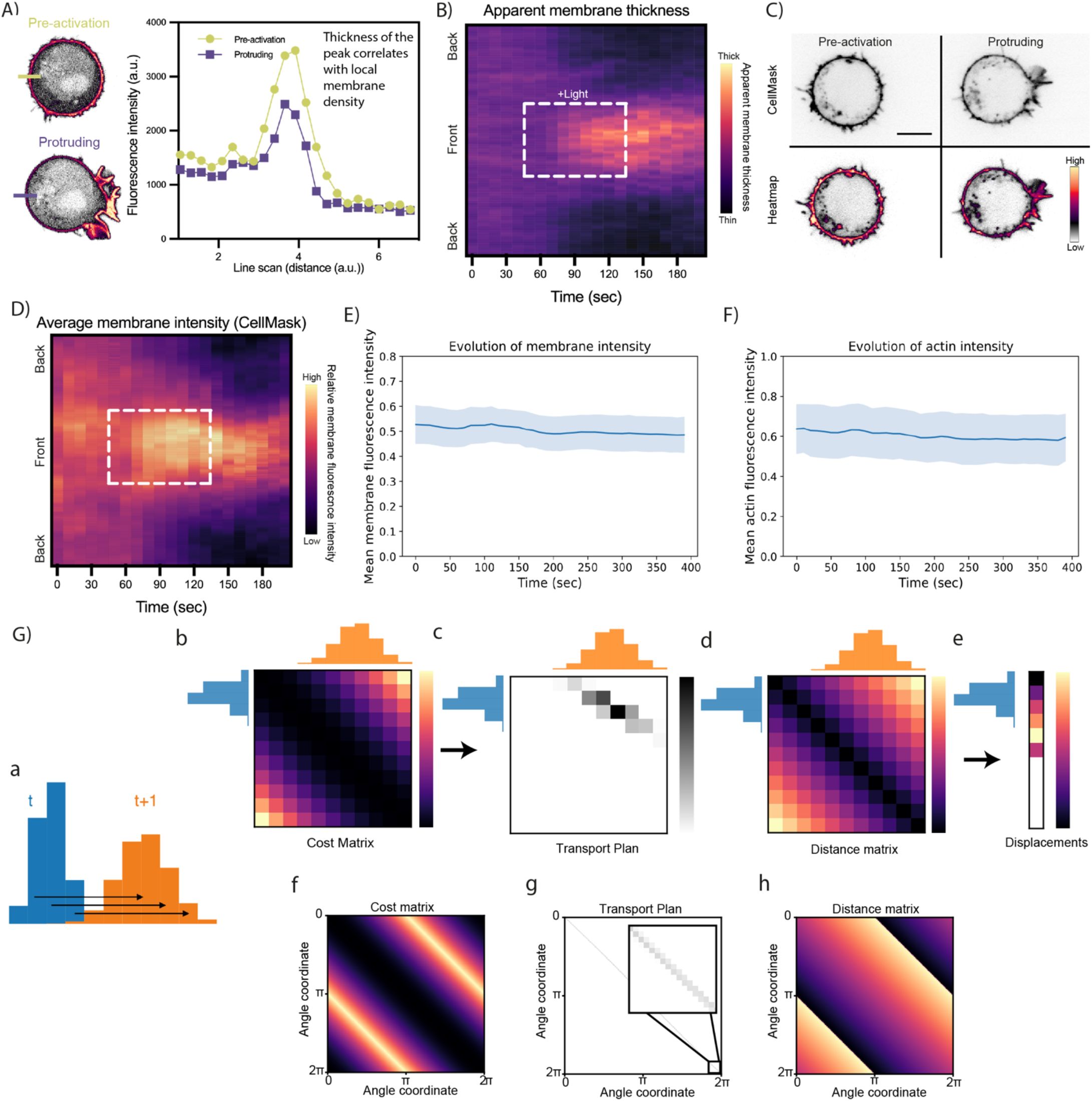
Long-range tension propagation coincides with directed membrane flows toward the protrusion. **(A)** Left: apparent membrane thickness is measured based on the width of fluorescence intensity profile across the cell contour, e.g., on the opposite side of cell protrusion (black line). At steady state (pre-activation) the cell membrane contour appears rugged (left image) and thick in width (light green curve in right plot), likely due to the presence of membrane reservoirs. As the cell protrudes, the membrane intensity outside of the protruding region drops (left image) and becomes thinner in width (purple curve in right plot). **(B)** Kymograph of averaged apparent membrane thickness along the normalized cell circumference (y-axis) over-time (x-axis): before, during, and after localized light-activated protrusion (box in white dashed line). On average, apparent membrane thickness reduces the most at the back of the cell (opposite from protruding front), likely reflecting a decrease in membrane reservoirs and a redistribution of extra membranes towards the protrusion site. **(C)** Representative confocal images of an opto-PI3K cell stained with plasma membrane dye (CellMask) before light activation or during protrusion. Scale bars: 5μm. **(D)** Kymograph of membrane fluorescence intensity (from cells stained with CellMask) along the normalized cell circumference (y-axis) over-time (x-axis): before, during, and after localized light-activated protrusion (box in white dashed line; n>25, N=4). (**E)** Evolution of the total membrane intensity across the cell contour. Except for a small intensity decrease due to the bleaching of the fluorophore, the membrane quantity is conserved. **(F)** Evolution of the total actin intensity across the cell contour. Bleaching of the fluorophore across time is visible. Actin intensity is conserved across time, with a higher standard deviation than the membrane intensity. **(G)** a-d) An illustrative example of optimal transport between two discrete 1-dimensional distributions, at time *t* (blue) and time *t+1* (orange), which represent the amounts of membrane (or actin) along the membrane contour at two different time points. a) Cost matrix C, where C[*i,j*] indicates the value of the cost to displace an element from position *i* to the position *j*. Here, the cost function shown is the square of the curvilinear distance. b) Transport Plan to go from the distribution at time *t* to the distribution at time *t+1*, minimizing the total cost of the displacement, computed from the cost matrix in a). c) Distance matrix D, where D[*i,j*] indicates the value of the distance between an element at the position *i* and an element at the position *j*. The distance chosen is the curvilinear distance. d) The transport plan and the distance matrix allow to compute the mean displacement for every position between times *t* and *t+1*. e-g) Matrices in the case of periodic boundary conditions, such as the circular contour of the cell. e) Cost matrix with periodic boundary conditions. The cost function chosen is still the square of the curvilinear distance, but as the topology of the curve is periodic, the matrix is changed to reflect this new topology. f) Example of the transport plan with real data. The matrix is almost diagonal, which means that the displacements reflect subtle changes of the distribution. g) Distance matrix with periodic boundary conditions. To keep track of the direction of the displacement, the distances can be positive or negative (and subsequently the positive and negative speed shown in Fig. 3F-H). A displacement in the clockwise direction (increasing angle coordinate) is positive, whereas a displacement in the counter-clockwise direction is negative.

**Fig. S7.**
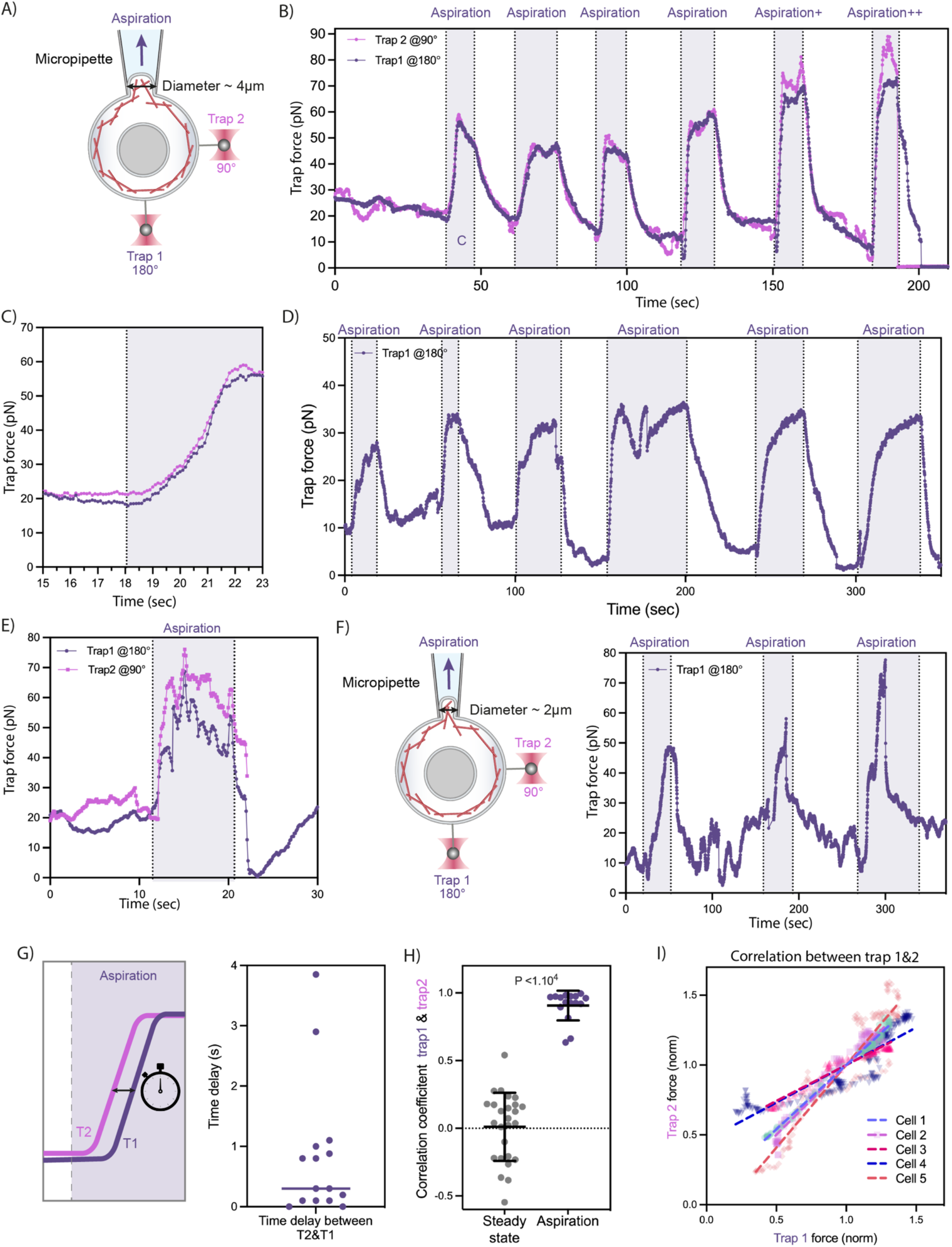
Mechanical perturbations applied on both membrane and cortex lead to rapid tension propagation across the cell. **(A)** A dual-tether pulling assay to simultaneously monitor membrane tension on the far-end (bottom, trap 1 at 180°) and on the side of the cell (right, trap 2 at 90°) during micropipette aspiration (top, ∼4-5 μm in tip diameter), which mechanically pulls on both the membrane and actin cortex underneath. (**B)** Representative time traces of dual trap forces over successive cycles of aspiration (shaded area) and relaxation; the magnitude of aspiration progressively increased in the last two cycles (+ and ++; the first three cycles were also shown in Fig. 4E). The nearly superimposable tension rise and fall on the two membrane tethers show that membrane tension propagates rapidly across the cell upon mechanical perturbations exerted to both the cortex and membrane. Note that the profiles of tension rise upon aspiration and of tension drop during relaxation resemble those observed with light-activated actin-driven protrusions (Fig. 2B). **(C)** Zoom-in on the first aspiration event shows that the trap force for membrane tension on the tether closer (pink) to the aspiration site started increasing slightly earlier and ended up slightly higher compared to that measured on the tether at the opposite cell pole from the protrusion (purple). **(D)** An example trace of tether tension response monitored on the opposite side of micropipette aspiration (trap 1 at 180°). Here, the recording lasted for six rounds of aspiration and relaxation. **(E)** Another example of dual-tether membrane tension measurement upon micropipette aspiration; the tether in trap 2 broke (*) shortly after the aspiration stopped. **(F)** Left: micropipettes of slightly smaller diameter (∼2μm) can also be used to exert mechanical suction (aspiration) on cells in the dual-tether pulling assay. Right: an example time trace of trap force for cell membrane tension exhibits robust responses over three aspiration cycles using a micropipette with ∼2-μm tip diameter. **(G)** Pearson correlation coefficient between dual trap forces measured before any perturbations (steady state) and during mechanical pulling upon micropipette aspiration. Error bar: means ± SD; p values from Welch’s unpaired Student’s t test (n>15, N>3). **(H)** Correlation plots of normalized trap forces between the two tethers during micropipette aspiration. Five representative measurements from different cells are shown; dashed lines: linear regression.

## Supplementary Text

### 1 Inference of surface flows with optimal transport

In this part, we derive the procedure allowing us to infer membrane and actin flows from time-varying kymographs of their fluorescence intensity at the cell surface.

#### 1.1 Kymographs generation

##### Raw kymographs

Our optogenetics procedure aims at inducing endogenous cell surface perturbations by illuminating light-excitable proteins in a given time window. These deformations were observed by staining the membrane or the actin cortex with a fluorescent marker (see Methods).

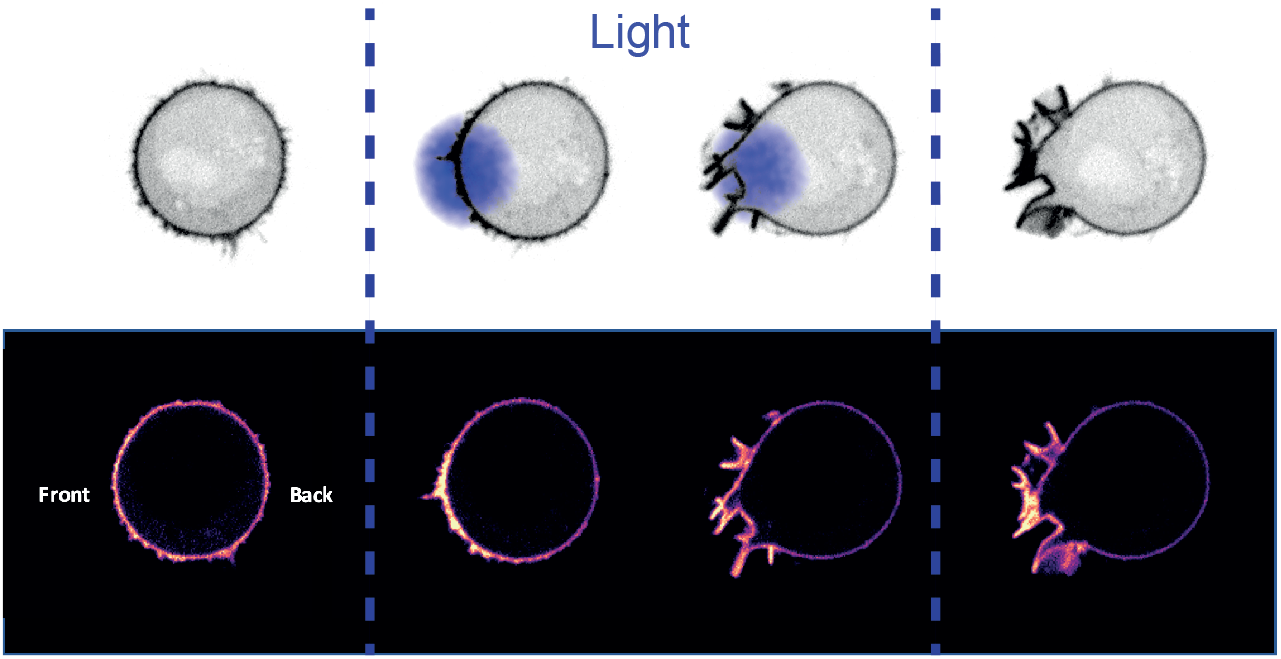

Normalized kymographs such as that shown in Fig.S6D are generated from 2D fluorescent timelapse images of membrane or actin by computing the distribution of membrane intensity across the contour of the cells. Images are first translated and rotated to have the center of the optogenetic activation zone always at the same point and the cell deformation in the same direction. Movies are segmented using cellpose, a deep-learning tool [1], then distributions of intensity are built by using an histogram where the *n* = 800 bins constituted regularly spaced angular sectors of the cells, centered around the centroid of the pixels of the segmented membrane.

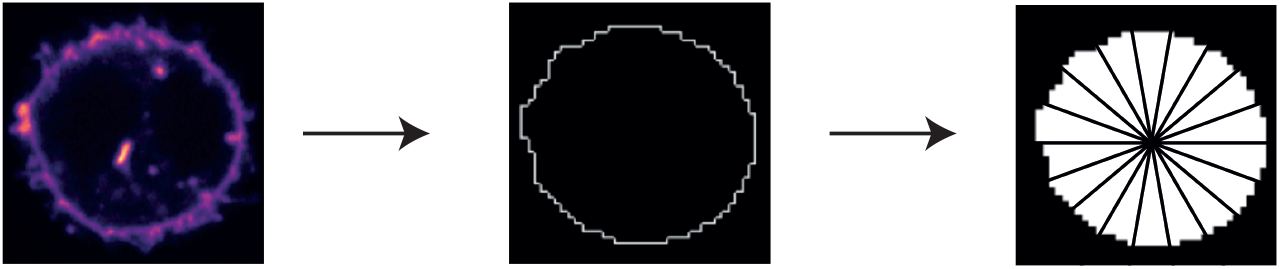
Microscopy images are segmented and membrane intensity is captured from the kymographs by binning the contour of the membrane

##### Total membrane and actin intensities are conserved in time

To observe membrane and actin flows, we imaged jointly membrane and actin during the optogenetics experiments (Fig. 3a,c of the main text). With the procedure introduced previously, we derived kymographs of membrane and actin evolution across time (Fig. 3b,d of the main text). When plotting the average membrane intensity, we observed that except for a bit of bleaching of the fluorophore, the intensity of the membrane signal is conserved across the time of one experiment (Fig. S6E). The same observation can be done for actin (Fig. S6F).

#### 1.2 Concise introduction on optimal transport

On the kymograph previously established, one may observe that at first, the distribution of membrane is uniform. After the illumination, the distribution becomes peaked at the front of the cell, becoming weaker at the back of the cell, before recovering to a uniform distribution after the illumination is stopped.

From this visual analysis, several questions arise:

- What kind of information can we extract from these observations?
- What kind of hypothesis do we have to make to infer a flow?

These questions can be tackled using transport theory, which is a mathematical field formalized by Gaspard Monge in 1781 [2], dedicated to the study of optimal transportation and allocation of resources. The problem was revisited in 1942 by Leonid Kantorovich who gave it its current formulation [3]. In our example, when we observe the distributions of membrane intensity at two consecutive timesteps *μ*_s_ and *μ*_t_, we observe that there has been a collective displacement of the membrane quantities between two positions. How to determine which membrane particle goes where? A first important physical hypothesis is that there is a notion of work W linked to each displacement: because of dissipation, a particle need to consume energy to realize a certain displacement through a path, that will be modeled by a cost function. Because of this cost, we expect that particles will undergo bounded displacements. Another hypothesis then is that displacements between two time-steps are as short as possible, i.e we assume that the time steps are sufficiently small to ensure that the two distributions are relatively close from each other and thus that between two timepoints individual particles are traveling using straight trajectories.

We choose a work that only depends on the length *l*_P_ of the path *P*, which is a good approximation only for small steps: 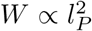. Since the time remains the same between two steps, this amounts to choose a cost function proportional to velocities squared: *W ∝ υ* ^2^, that is akin to a friction-like dissipation cost function. This is physically a fair hypothesis for inferring the flows of membrane, as far as we suppose in this manuscript that they are limited by the relative friction to the cortex (see model description in Section 2 of this Supplementary Text).

We can reformulate the flow inference as an optimisation problem. We first introduce the distance function *d*(·, ·), that takes as arguments spatial positions along the membrane and returns a scalar value that represents the distance between two points. We write *c*(·, ·) the cost function that takes as arguments spatial positions along the membrane and returns a scalar value that represents the cost to go from one place to another. In the following, we will use *c*(·,·) = *d*^2^(·,·). We call the transport *m* a map such that *m*(*μ*_*s*_) = *μ*_*t*_. We want to find the optimal transport *m*^*^ that minimizes:

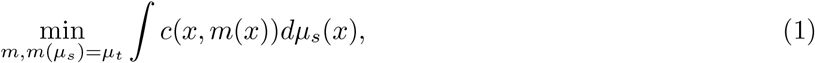

where the integration is done on the probability measure *μ*_s_(*x*).

The optimal cost *W*_*d*_(*μ*_*s*_, *μ*_*t*_) = ∫ *c*(*x, m*^*^ (*x*))*dμ*_*s*_(*x*) is called the Wasserstein distance between two measures. Here we are more interested on the mapping on itself, that is called the optimal transport matrix. Indeed, as this matrix gives a notion of distance traveled at each point on the membrane, we will determine the speed of the membrane flows using these discrete displacements.

#### 1.3 Discrete periodic optimal transport

Here we study discrete distributions. Following the notations of [4], we define a cost matrix C, a transport matrix M and a distance matrix D, (Fig S6G.a-c), discretized versions of their continuous surrogates *d*(·, ·) *c*(·, ·), and *m*(·). For each discrete bin i and j, of positions *x*_*i*_ and *x*_*j*_, we have : *D*_*i,j*_ = *d*(*x*_*i*_, *x*_*j*_). As well, 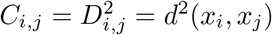. We write 𝟙 the column vector where each value is one. For two simple distributions a and b, the optimal transport *M*^*^ is defined by:

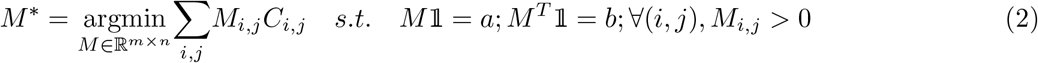

We compute the optimal transport *M*^*^ using the standard Earth Mover’s Distance (EMD) with [4]. This exact computation does not scales well with the number of bins (It has a *O*(*n*^3^) complexity [5]) but the 1D case has a low enough number of points to obtain results in a reasonable amount of time.

The velocity field V is obtained by averaging the distance traveled for every element, and dividing by the timestep *δt* (Fig S6G.d):

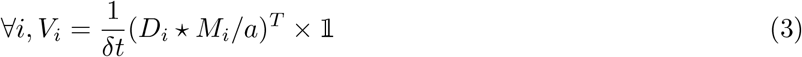

Where ⋆ is an element-wise multiplication, / the element wise division and × is the standard matrix product. *X*_*i*_ denotes the *i*th column of the matrix *X*_*i,j*_.

##### Optimal transport with periodic boundary conditions

In our use-case, the cells are circular, and thus our distributions have periodic boundary conditions. One can travel along the membrane using a clockwise and an anticlockwise path. To take this into account, we adapted cost and distances matrices (Fig. S6G.e,g). Moreover, we decided to compute algebraic distances (i.e positive or negative values), to determine the direction of the flow from its sign (Fig. 3F,H).

#### 1.4 Applications to neutrophil-like HL60 cells

##### 1.4.1 Lipid membrane flows

For each cell, we generated an individual kymograph, from which we performed the flow inference using optimal transport (see image bellow). Velocity fields were then averaged across all our movies (Fig. 3F).

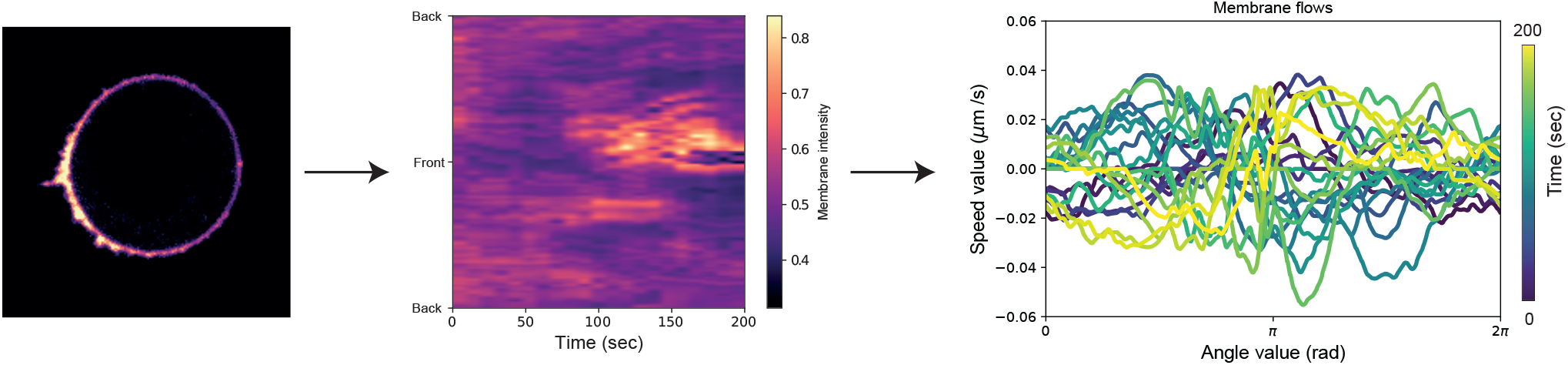
The movies are used to compute kymographs that are used to compute velocity fields

We proposed an alternative representation of the flow (Fig. 3G) by representing it with arrows of length that scales with the intensity of the displacement measured (i.e with the speed of the flow): anterograde flows (from back to front) were displayed in red while retrograde flows were displayed in blue. We see clearly three phases: t=20 sec: Before illumination, nothing happens, there is a residual noise that is averaged across all the samples, which make the velocity really small. t=70sec During and Right After illumination: there is a anterograde flows, very fast and very intense. t=170sec. A retrograde flow, more progressive, restores the initial configuration of a uniformly distribution of membrane.

We provide a ([Jupyter Notebook](https://github.com/VirtualEmbryo/membrane-cortex-tension.git/Membrane_flows_notebook.ipynb)) and the original data for reproducibility on [Github](https://github.com/VirtualEmbryo/membrane-cortex-tension).

##### 1.4.2 Cortical actin flows

We reproduce this procedure with actin. We generated kymographs of actin for each cells, and velocity fields from these kymographs. Velocity fields are then averaged across cells (Fig. 3H). We see the same behavior than with membrane flows: anterograde and then retrograde flows (Fig. 3I). We provide a ([Jupyter Notebook](https://github.com/VirtualEmbryo/membrane-cortex-tension.git/Actin_flows_notebook.ipynb)) and the original data for reproducibility on [Github](https://github.com/VirtualEmbryo/membrane-cortex-tension).

## 2 Composite membrane-cortex mechanical model

### 2.1 Model hypotheses

We model the cell surface as a composite structure composed of a viscous active actomyosin cortex linked to an elastic membrane by adhesion proteins anchored in the cortex, as represented on the sketch below. The force transmission between membrane and cortex is mediated by a friction term. Our model hypotheses are directly inspired by the picket fence theory of the plasma membrane [6].

To simplify the description of the cortex-membrane interaction and exploiting the axisymmetry of the problem, we derive a minimal one-dimensional model of the composite cortex-membrane cell surface. The cortex-membrane structure is a straight line with spatial coordinate *ξ* ∈ [0, *L*], where *L* is the curvilinear length of the structure. We derive our model in a discrete setting^1^, where we subdivide the structure into *N* elements of size *L*/*N*, labelled by an index *i*.

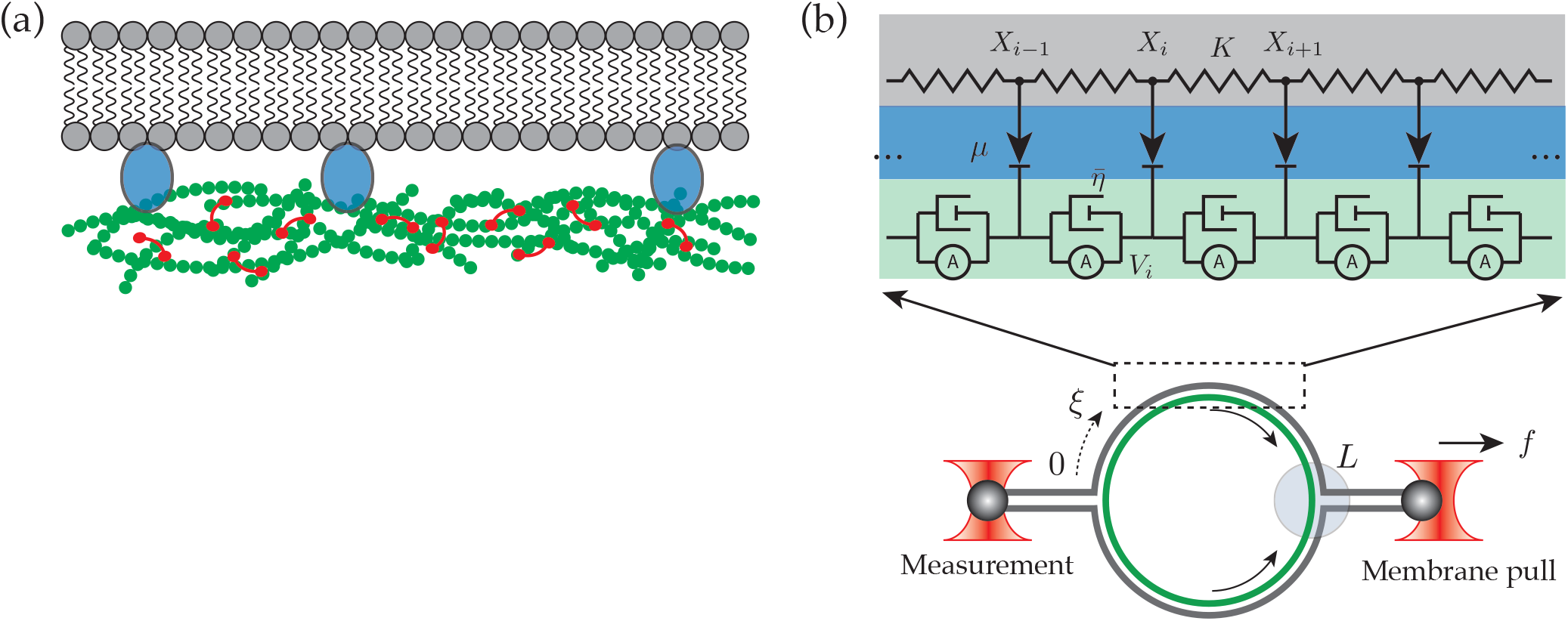
Sketch of the composite membrane-cortex structure and its mechanical representation. (a) Membrane-cortex structure. The plasma membrane is a passive bilayer that is attached to the cortex via adhesive molecules (blue ellipses) that are firmly attached to the cortex but can tangentially flow along the bilayer. (b) Schematic representation of the cortex-membrane structure. Assuming a axisymmetric geometry of the system, we can use a one-dimensional model for the cell surface. Multiple optical traps are used for both application and measurement of force. Cortical flows (arrows) may arise from localized light-based protein activation.

The discrete model of the membrane consists of a chain of elastic springs with elastic constant *K*[*N*/*m*]. To each spring *i* it is associated a displacement field *X*_*i*_. The actomyosin cortex is also represented here in its discrete form using a phenomenological model for the contractility, in which the rheology of the cortex is equivalent to the parallel association of a contractile element and a dashpot, representing the viscous dissipation [7]. Such an element is characterized by an active tension *σ*_*i*_, a bulk viscosity *η*_3*D*_ and a velocity field *V*_*i*_. The membrane-cortex interaction is ensured by the presence of cortex-bound proteins that are linked to the plasma membrane but allow for relative motion to the cortex, which generates frictional forces proportional to the relative velocity between the membrane and cortex and characterized by a friction coefficient 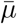.

### 2.2 Mechanical equations of the composite membrane

The system of equations that govern the dynamics of this system derives from force balance in both the cortex and the membrane.

#### 2.2.1 Discrete equilibrium equations

To obtain the mechanical equilibrium at a position *X*_*i*_ in the membrane we consider the force balance between the elastic response of the membrane and the friction force arising from cortex-membrane interaction. The discrete equation at a node *i* reads

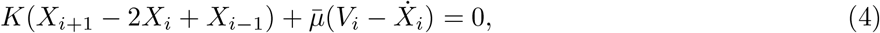

where 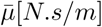 is the friction coefficient of the cortex-membrane interaction, *V*_*i*_ is the velocity field of the cortex at node *i*, and 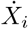 is the time-derivative of the membrane’s displacement field.

Analogously, we can write down the equilibrium for an element *i* in the cortex. Let *η*_3*D*_[*Pa*.*s*] be the three-dimensional effective viscosity of the actomyosin gel of constant thickness *T*_0_, and let define a characteristic cross area element *δ A* = *T*_0_*h*, where h is a characteristic transversal length. The force due to the viscous deformation of the cortex is *δAη*_3D_ ∂ *V*_*i*_/*∂ξ*. The balance of forces at the element *i* reads then

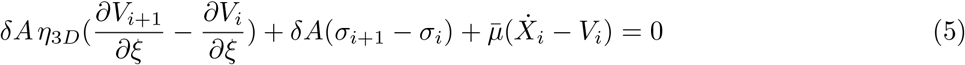

#### 2.2.2 Continuum limit

In the continuum limit, we substitute the discrete derivatives *X*_*i*+1_ − *X*_i_ = *h* ∂*x*(*ξ*)/*∂ξ*, where *h* = *L*/*N* is the discrete spacing and tends to zero as *N* → ∞. Let 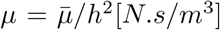 represent the local friction density. For notation uniformity, the elastic constant *K* is written as *k ≡ K*[*N*/*m*]. We can write a continuum analog of (4) as,

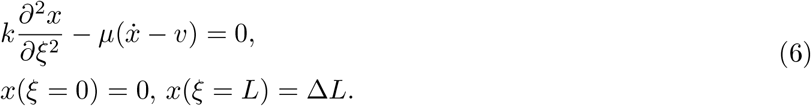

Notice that we enforce a Dirichlet boundary condition on the displacement at *ξ* = 0 such that the displacement field is zero to ensure the symmetry of the model. We further suppose that the mechanical perturbation applied by the optical trap can be modelled by an additional Dirichlet boundary condition at *ξ* = *L*.

The continuum analog of (5) is

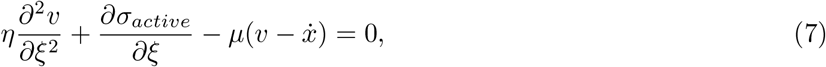

where *η* = *T*_0_*η*_3*D*_[Pa.s.m] is the two-dimensional effective viscosity of the cortex^2^.

The system can be seen as a one-dimensional representation of a axisymmetric system. It is easy to visualise that if a localized contraction happens somewhere in the cortex, there will be always a point in the cell in which the cortical flows are identically zero because of symmetry. Our one-dimensional representation is then seen as the “unwrapping” of such system into a one-dimensional string, where we enforce the boundary conditions *υ* (ξ = 0) = 0, and 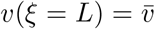, where 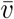 is the velocity of protuding membrane at the cell front in response to light activation.

#### 2.2.3 Final system of mechanical equations

The cortex-membrane system is represented by a coarse-grained model in which the membrane state is fully characterized by the scalar *displacement* field *x*, while the cortex is described by the scalar *velocity* field v. The one-dimensional spatial domain is Ω ∈ [0, *L*], and we observe the sytem at a time *t* ∈ (0,*T*]. Dividing both equations (6) and (7) by *μ*, the final system of equations describing the system dynamics reads

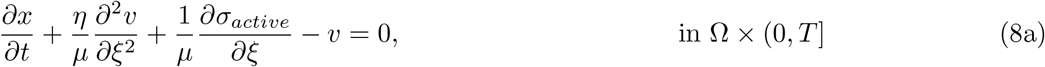

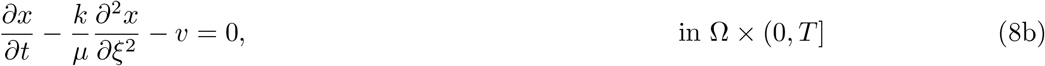

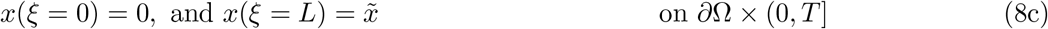

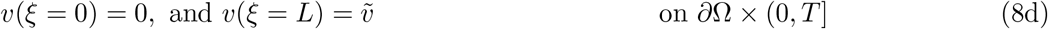

One may identify three main parameters controlling the system dynamics:

- A hydrodynamic length 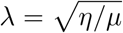, which compares membrane friction and cortex viscous forces and corresponds to the typical length above which cortical flows may be dampened by membrane-cortex friction, and to be compared with the system lenght *L*.
- A characteristic timescale *τ* = *η*/*k*, which compares the elastic membrane force with the cortical viscous force.
- A dimensionless cortex pulling velocity 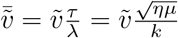.

Note that, only spatial gradients, but not the absolute value of the cortical tension *γ* = *T*_0_*σ*_*active*_ does play a role in the system response dynamics.

These length and time-scales could allow us to non-dimensionalize the system above in the following form

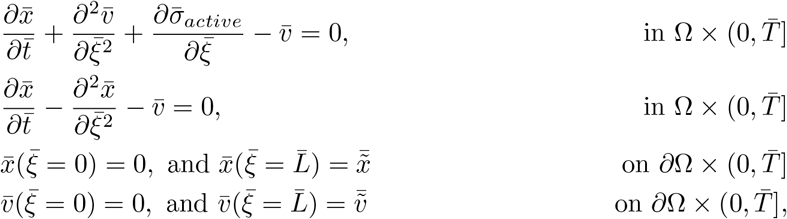

We decided yet to keep dimensionalized equations in the next for easier physical interpretation.

### 2.3 Numerical implementation

Because we have a time-dependent system of partial differential equations (PDEs), we first discretize in time by a finite difference approximation, which yields a sequence of stationary problems, and then turn each stationary problem into a variational formulation. From now on, the derivative with respect to the spatial coordinate *ξ* will be denoted by ∇ ≡ *∂*/ *∂ξ*.

#### 2.3.1 Time discretization

In order to solve (9), for the displacement *x*(*ξ, t*) and velocity *υ* (*ξ, t*), we first discuss the time discretization. We use the superscript n to denote a quantity at discrete timesteps *t*^*n*^ = *n*Δ*t*, where *n* = 0, 1, 2,. For an arbitrary field *ϕ* (*ξ*, t), *ϕ*^*n*^ = *ϕ* (*ξ,t* = *t*^*n*^). A finite difference implicit discretization in time first consists of sampling the PDE system at some time, say *t*^*n*+1^:

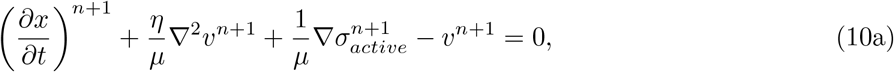

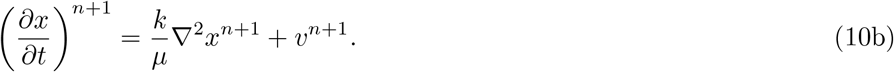

The time-derivative is then approximated by a difference quotient. For simplicity and stability reasons, we chose a simple backward Euler method:

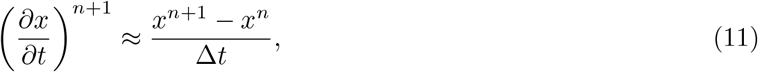

where Δ*t* is the time discretization parameter.

#### 2.3.2 Variational formulation

Let 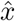, and 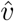 be the test functions. We first approximate the time derivatives using (11), then mutiply each equation by tests function, integrate over the spatial domain Ω, and sum the equations.

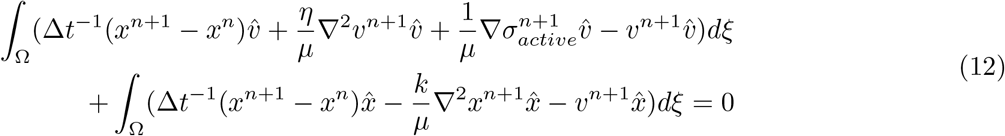

Integration by parts allows us to write,

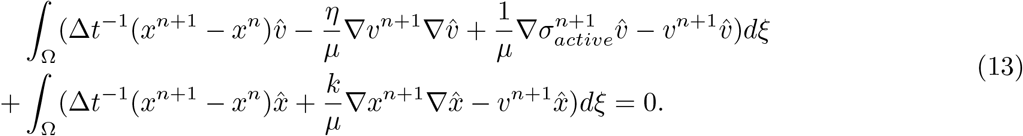

#### 2.3.3 Finite-element implementation

At each time step, we solve (13) for the cortex velocity field *υ*, and membrane displacement *x*. We assume a quadratic continuous (P2-Lagrange finite-element) interpolation for the velocity field *υ*, and *x*. The dynamic evolution is solved using a backward Euler method.

Aiming at facilitating the spread, modification, refinement of our model and to highlight its features, we furthermore devise a numerical scheme (available on GitHub) to solve the system of partial differential equations with the finite-element library FEniCS [8].

#### 2.3.4 Control case: constant contractility

As a validation for our finite element implementation of the coupled system of equations and boundary conditions, we compare the results of our code against a simplified setting where analytical solutions are possible. The simplest case assumes a uniform contractile stress, and we set *υ* = 0 to solve the membrane equilibrium equation

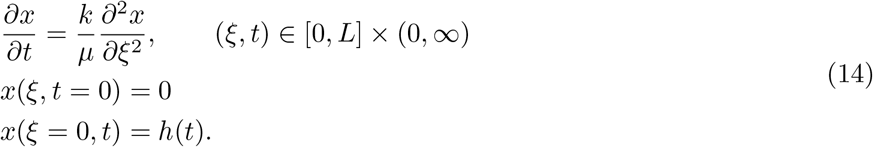

which is similar to a heat equation with boundary conditions. Such equation has a closed analytical solution

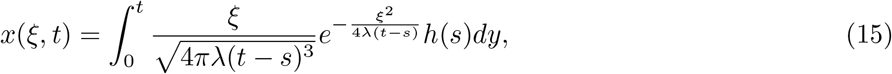

where *k*/*μ* = λ.

Numerically, we need to decouple the membrane equilibrium equations from the cortex. This is achieved by setting *μ* → 0. Because (14) only depends on the ratio *k*/*μ*, we make the limit *μ →* 0, and *k* → 0, such that *k*/*μ* → 1. The comparison between analytical results and finite element simulations shows good agreement, as shown below

### 2.4 Numerical values

The physical values used in the simulations are summarized in the table below. The membrane elasticity [N/m] is estimated based on the membrane tension, which can be obtained by pulling membrane tethers from the plasma membrane using laser tweezers [9]. The friction coefficient here is estimated as *μ* = *ζ ρ*_0_, where *ζ* ≈ 10^−^6 Pa.s.m [10] is the drag resistance of an individual linker and *ρ* = 10^14^*m*^−2^ is the surface density of linkers [11].

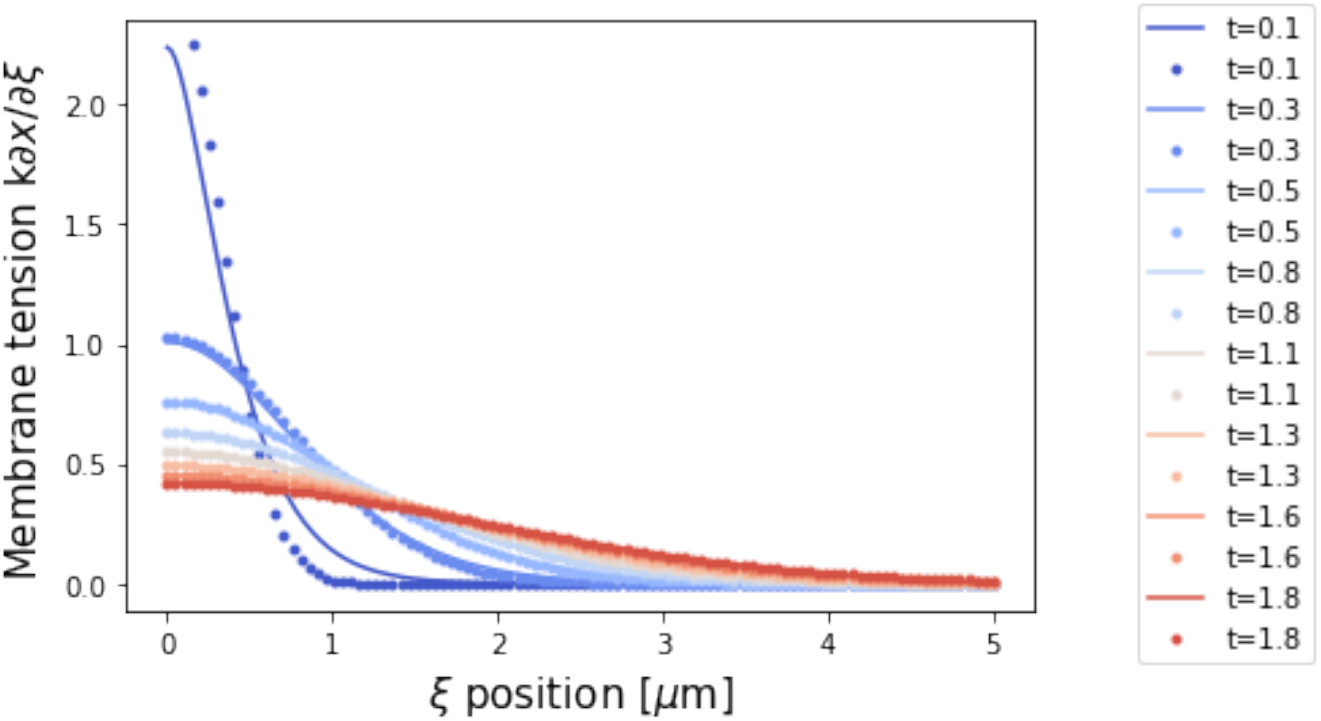
Membrane tension profiles along the membrane for increasing times. *Continuous line: analytical solution based on* (15). *Points: finite element solution of* (13).

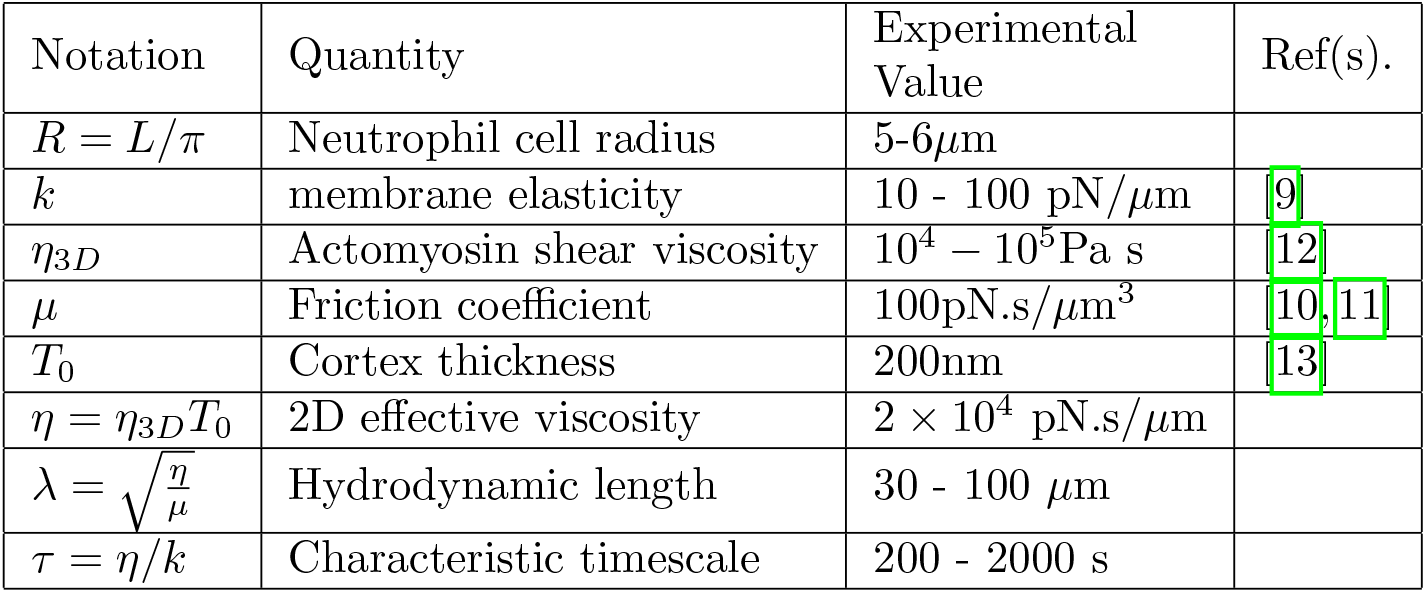
Table of numerical values.

### 2.5 Results

We provide a ([Jupyter Notebook](https://github.com/VirtualEmbryo/membrane-cortex-tension.git/main.ipynb)) with the code and implementation for figures in the main text (Figs. 4 B-C) and figures for the control above and for the role of friction (see below).

In the article, we are mainly interested in understanding how the membrane (i.e. lipid bilayer) tension propagates along the cell. There are two main cases of interest. In the “exogenous perturbation”, the plasma membrane is pulled by an optical trap at the cell front and a membrane force is measured by an optical tweezer at the other end (back of the cell). In the so-called “endogenous perturbation”, we used an optogenetic tool to locally generate and actin-driven cellular protrusion of velocity 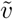 at the cell front and we measure the instantaneous resulting membrane tension at the back. The local membrane tension measured by an optical trap may be defined as

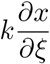

#### Exogenous perturbation

We modelled the perturbation on the membrane by an optical trap as a Dirichlet boundary condition on the displacement field *x*(*ξ* = *L, t*). The perturbation propagates along the cell because of the elasticity of the membrane, however, the viscous dissipation of the membrane-cortex interaction buffers such propagation. It is important to notice that the observational timescale (The time at which the membrane tension is observed) is crucial. Since the membrane is considered to be linearly elastic, in the limit *t* → ∞, the tension becomes uniform across the whole cell. Here we chose to solve the system up to some observational time *t*_*f*_. In the figure below, we illustrate a scheme for the exogenous perturbation, where a cell is pulled at one end and we observe the force propagation along the system.

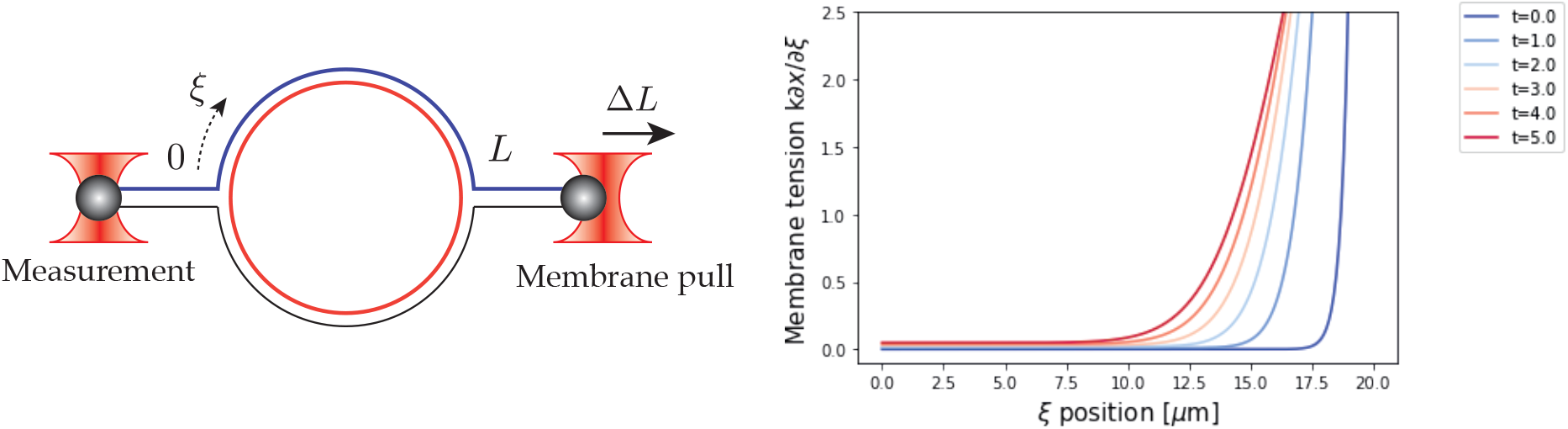
Exogenous perturbation. Sketch of the system (left) and membrane tension profiles [nN] for increasing times after the perturbation (right). Refer to [GitHub](https://github.com/VirtualEmbryo/membrane-cortex-tension.git/main.ipynb) for the set of physical parameters and procedure.

#### Endogenous perturbation

The endogenous force generation can be modelled in two different ways providing same qualitative features. First, as an active source of contraction via the term *σ*, or via a Dirichlet boundary condition on the velocity field 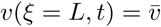, which is used here for simplicity. The boundary condition is complemented with a full clamp at *ξ* = 0, and by unconstraining the membrane at *ξ* = *L*. As in the endogenous case, here we need to specify an observational time, which is linked to the fact that the contractility is happening at a definite, finite, time interval.

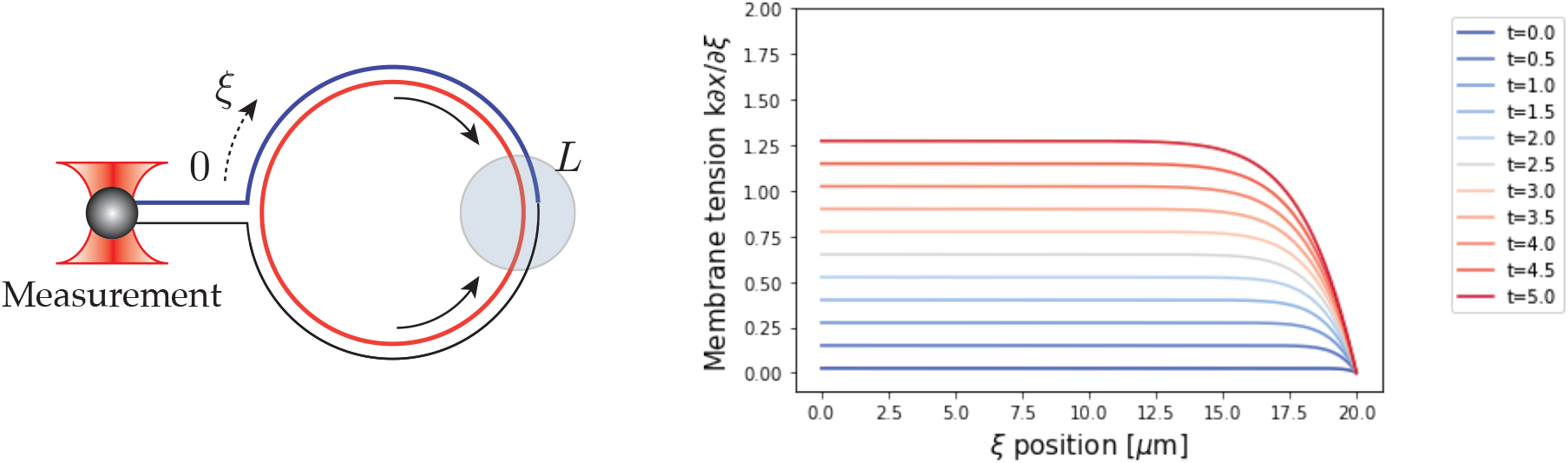
Endogenous perturbation. Sketch of the system (left) and membrane tension profiles [nN] for increasing times after the perturbation (right). Refer to GitHub for the set of physical parameters and procedure.

#### The role of friction

A key factor in the system is the role of the cortex-membrane friction, that is proportional to the relative particle velocity between the cortex and membrane and is expected to be proportional to the surface density of membrane-cortex linkers.

In the case of an **exogenous perturbation**, a perturbation in the membrane will propagate as in a elastic solid in the limit of low friction, *μ* → 0, and the membrane tension will be uniform across the cell. In the high friction limit *μ* ≫ 1, the perturbation is dissipated along the cell and the force is not propagated at finite observational time, as shown on panel (a) of the figure below. Furthermore, the cortex viscosity plays here an important role in dampening the tension propagation, as illustrated in the panel (b) of the figure below.

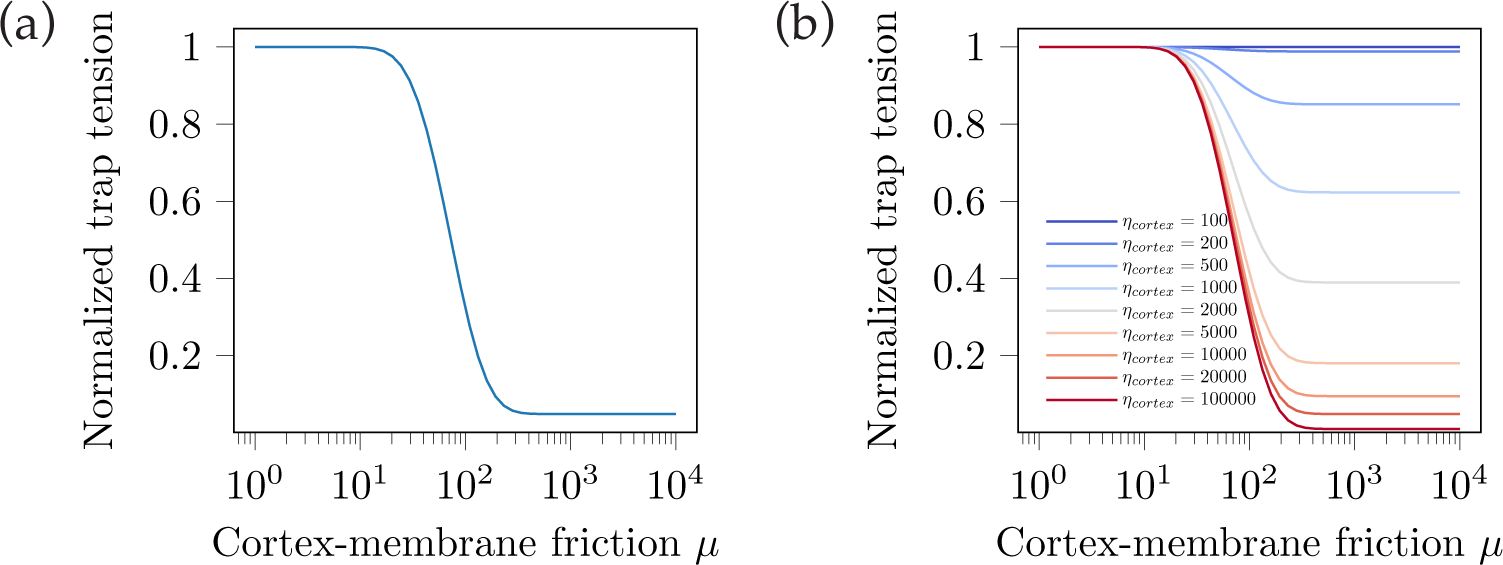
Normalized membrane tension for an exogenous perturbation as a function of the cortex-membrane friction. (*a) Normalized membrane tension profile for a cortex viscosity η* = 100000 *pN*.*s/*mu*m*. (*b) Normalized membrane tension profiles for increasing cortical viscosities η* = 10^2^ − 10^5^. *Refer to [GitHub](https://github.com/VirtualEmbryo/membrane-cortex-tension.git/main.ipynb) for the whole set of physical parameters and procedure*. The tension is normalized by its max value (tension = tension/max tension).

In the case of an **endogenous perturbation**, force transmission behaves depends on the cortex-membrane friction in a different manner. In the low friction *μ* → 0, the cortex does not interact with the membrane, meaning that cortical flows will not generate membrane stresses. In the high friction limit *μ* » 1, cortical flows will drag the membrane transmitting the tension, as shown on the figure below.

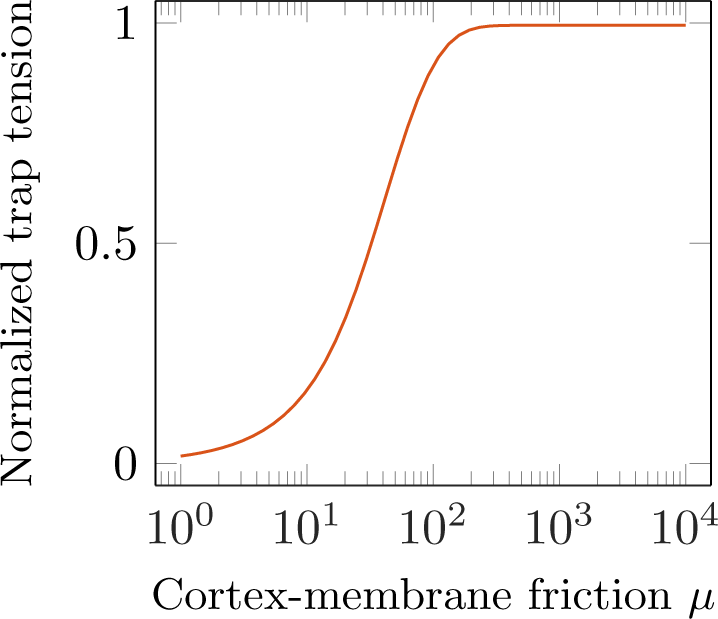
Normalized membrane tension for an endogenous perturbation as a function of the cortex-membrane friction. Refer to GitHub for the set of physical parameters and procedure. The tension is normalized by its max value.

## Footnotes

1 Note that both cortex and membrane share the same spatial coordinates. In a discrete setting, this means that a particular spatial point will correspond to the same index *i* for both cortex and membrane.

2 Notice that we use lowercase letters *x, v* when describing continuum variables, while uppercase *X*_*i*_, *V*_*i*_ will be reserved for discrete variables

